# Heat shock protein 90 is a master regulator of HIV-1 latency

**DOI:** 10.1101/2024.08.23.609311

**Authors:** Somaya Noorsaeed, Nawal AlBurtamani, Ahmed Rokan, Ariberto Fassati

## Abstract

An estimated 32 million people live with HIV-1 globally. Combined antiretroviral therapy suppresses viral replication but therapy interruption results in viral rebound from a latent reservoir mainly found in memory CD4+ T cells. Treatment is therefore lifelong and not curative. Eradication of this viral reservoir requires heterologous ΔCCR5 hematopoietic stem cell transplantation, which is not broadly applicable. Alternative cure strategies include the pharmacological reactivation of latently infected cells to promote their immune-mediated clearance, or the induction of deep latency. HIV-1 latency is multifactorial and linked to the activation status of the infected CD4+ T cell. Hence to perturb latency, multiple pathways need to be simultaneously targeted without affecting CD4+ T cell function. Hsp90 has been shown to regulate HIV-1 latency, although knowledge on the pathways is limited. Because hsp90 promotes the proper folding of numerous cellular proteins required for HIV-1 gene expression, we hypothesized that hsp90 might be a master regulator of latency. We tested this hypothesis using a polyclonal Jurkat cell model of latency and ex-vivo latently infected primary CD4+ T cells. We found that hsp90 is required for HIV-1 reactivation mediated by the T-cell receptor, phorbol esters, TNF-α, inhibition of FOXO-1, and agonists of TLR-7 and TLR-8. Inhibition of hsp90 abrogated activation of the NF-kB, NFAT and AP-1 signal transduction pathways, and this phenotype was recapitulated by targeting TAK1, an hsp90 client protein. Within the CD4+ T cell population, naïve and effector memory cells were most sensitive to hsp90 inhibition, which did not perturb their phenotype or activation state. Our results indicate that hsp90 is a master regulator of HIV-1 latency that can potentially be targeted in cure strategies.

**Author summary:** HIV-1 affects around 32 million people globally. Current treatments, known as combined antiretroviral therapy, can suppress the virus but do not cure the infection and if the treatment stops, the virus comes back. This happens because the virus hides in a population of immune cells called memory CD4+ T cells. To truly cure HIV-1, some strategies involve complex and risky procedures like hematopoietic stem cell transplants, which are not widely applicable. Another approach is to reactivate the hidden virus in the cells, so the immune system can eliminate it, or to force the virus into an even deeper hiding state. HIV-1 latency, or its ability to hide in cells, is influenced by many factors and cells need to be activated to disrupt it. Hsp90 is a chaperone that regulates the function of numerous proteins important for HIV-1 latency and is known to play a role in maintaining this hidden state of the virus. We therefore wondered if Hsp90 acts like a master regulator of latency. Using lab-based models, we discovered that Hsp90 is crucial for the reactivation of HIV-1 through various pathways. By inhibiting Hsp90, the activation of key signalling pathways necessary for viral reactivation was blocked. Importantly, blocking Hsp90 did not harm the CD4+ T cells’ function or state. Hsp90 inhibitors, already tested in cancer treatments, could thus be a promising avenue for HIV-1 cure strategies, as they seem to hold the key to maintaining HIV-1 latency.

## Introduction

Untreated human immunodeficiency virus type 1 (HIV-1) infection causes a progressive loss of CD4+ T cells that eventually results in death of the infected individual due to profound immunodeficiency, opportunistic infections, neurocognitive deficits and cancer such as Kaposi sarcoma [1, 2]. Currently, combined antiretroviral therapy (cART) extends the life expectancy of people living with HIV-1 (PLWH) who are virologically suppressed to levels almost equal to uninfected individuals [3]. Treatment with cART must be lifelong because its interruption results in rapid virological rebound from a latently infected cell reservoir, even if therapy started early post-infection [4]. The largest reservoir is constituted by memory CD4+ T cells in which HIV-1 has integrated and established a transcriptionally silent infection, although infected tissue resident macrophages and microglial cells in the brain also contribute to the reservoir [5, 6]. The latent reservoir is long-lived due to homeostatic proliferation of latently infected memory CD4+ T cells, their clonal expansion and immune selection, and the ability of intact but silent HIV-1 proviruses to reactivate upon stimulation, restarting the replication cycle [5, 7]. Thus, cART cannot cure HIV-1 infection. To compound this problem, during cART, many PLWH show persistent immune activation due to gut microbial translocation, co-infections and viral antigen expression from a proportion of both intact and defective HIV-1 proviruses [8]. Although cART has mostly eliminated AIDS-defining illnesses, persistent immune activation significantly increases the odds developing co-morbidities in PLWH [9].

A cure by eradicating HIV-1 infected cells has been achieved only by allogeneic hematopoietic stem cell transplantation using homozygous ΔCCR5 donors, but this procedure is not broadly applicable to PLWH [10]. However, approaches to achieve long-term remission after therapy interruption are being developed [11]. One approach combines latency reactivating agents (LRAs) to stimulate viral antigen expression followed by immune-mediated clearance of infected cells. Another approach aims at repressing HIV-1 reactivation and generating a state of deep, potentially irreversible latency [11]. For optimal results, these two strategies require a detailed understanding of how HIV-1 latency is regulated and maintained.

HIV-1 gene expression is driven by the enhancer and promoter regions in the viral long terminal repeats (LTRs). Initiation of HIV-1 transcription relies on the transcription factors NFAT, NF-kB, AP-1 (c-Fos/c-Jun) and the ras-responsive binding factor-2 (RBF-2), which bind to cognate cis-acting sequences in the LTR [12–14]. Notably, these transcription factors determine not just HIV-1 transcriptional initiation, but also many of the cellular response to T-cell receptor (TCR) stimulation, demonstrating the close link between T-cell and viral transcriptional regulation [15–17]. Upon transcriptional initiation, the viral protein Tat is produced and imported into the nucleus where it recruits the host positive transcription elongator factor P-TEFb on to the LTR [18, 19]. P-TEFb, a heterodimer complex formed by cyclin T1 (CyT1) and the kinase subunit CDK9, boosts processivity of RNA polymerase (RNAPol) II by phosphorylating its C-terminal domain and triggering the release of negative regulators such as DSIF and NELF [18, 19]. This is followed by the epigenetic reorganization of chromatin at the site of viral integration, leading to sustained viral gene expression [18, 19].

HIV-1 latency is established at multiple levels, including the integration site in the host genome, the epigenetic configuration of chromatin at or near the site of integration, the activation state of the infected CD4+ T cells, which determines the nuclear availability of critical transcription factors, the abundance of P-TEFb, and the recruitment of transcriptional and post-transcriptional repressive factors [18, 19].

Inhibiting HIV-1 gene expression should prevent HIV-1 reactivation from latency and reduce chronic inflammation in PLWH, yet there are no approved drugs that target this step of the viral life cycle, and only a few are at the development or pre-clinical stage. Examples include PKC, PI3K and MEK inhibitors, which have been reported to suppress HIV-1 latency reactivation *in vitro* by inhibiting TCR mediated stimulation of latently infected CD4+ T cells [13, 20]. Aryl hydrocarbon receptor (AhR) agonists [21] and RORC2 agonists have been reported by our group to block viral outgrowth after stimulation of patients’ latently infected CD4+ T cells [22]. One limitation of this approach is that HIV-1 reactivation from latency depends on several parallel signalling pathways [19] hence blocking one pathway at a time may not be sufficient to achieve measurable silencing of viral gene expression. In addition, the effect of these drugs on CD4+ T cell behaviour is poorly understood.

This limitation can be addressed by targeting a viral protein that is essential for HIV-1 gene expression. An analogue of cortistatin A has been shown to suppress Tat activity both *in vitro* and in humanized mice, resulting in specific inhibition of HIV-1 reactivation, and fostering epigenetic modifications that maintain latency longer term [23–25]. A complementary strategy is to target master regulators that control multiple parallel pathways required for HIV-1 reactivation. Thus, there is a clear rationale to develop pharmacological interventions that repress HIV-1 reactivation from latency by targeting multiple pathways simultaneously, provided that these interventions do not perturb the overall T cell function.

We and others have demonstrated that heat shock protein 90 (hsp90) is a host factor required for HIV-1 gene expression and reactivation from latency [13, 26–33]. The chaperone seems to act mainly at the early stages of viral gene expression [26, 27, 32, 34]. Hsp90 and its co-chaperone Cdc37 were found to promote the activation of the NF-kB pathway by ensuring the correct assembly and function of the IKK complex [13]. Hsp90 also co-localized with actively transcribing proviruses [13] and maintained the function of P-TEFb [32, 34]. Notably, treatment of latently infected cells with selective Hsp90 inhibitors *in vitro* and in humanized mice repressed HIV-1 reactivation, and this effect was long-lasting [13, 28, 31, 32, 34]. Several selective hsp90 inhibitors have been tested in phase II and III clinical trials to treat haematological malignancies and solid tumours [35]. The pharmacokinetics and pharmacodynamics of these drugs are well understood, and the drug concentration sufficient to inhibit HIV-1 reactivation is significantly lower than that one required for the anti-cancer effect [13, 28, 35, 36], paving the way to their possible repurposing to inhibit HIV-1 reactivation in PLWH.

Hsp90 is a very conserved molecular chaperone that is responsible for folding, stabilizing, and activating a broad variety of client proteins. It operates as part of a large protein complex that includes co-chaperones, such as hsp70, and other regulatory factors that control the activity and stability of client proteins. Hsp90 is known to facilitate many cellular processes, including protein homeostasis, signal transduction, and stress response [37]. Because hsp90 promotes the correct folding and assembly of many host factors required for HIV-1 gene expression [13, 27, 28, 31, 34, 38], we hypothesized that hsp90 might be a master regulator of HIV-1 reactivation from latency. Here, we tested this hypothesis using a polyclonal Jurkat cell model of latency and ex-vivo latently infected primary CD4+ T cells, and we evaluated how inhibiting hsp90 affects the activation and differentiation of CD4+ T cells.

## Results

### Establishment of a polyclonal Jurkat model of HIV-1 latency

To investigate if hsp90 is a master regulator of HIV-1 latency, we first established a cell model of latency in which a variety of latency reversing agents (LRAs) and their dependence on hsp90 could be easily tested. The J-Lat cell clones have been very useful to study the regulation of HIV-1 latency and many of the key findings that emerged using this model have been confirmed in primary CD4+ T cells and in cells from PLWH [39–42]. However, the clonality of the J-Lat model, in which each clone has a single provirus integrated at a specific location, is a limitation. Because the site of integration affects HIV-1 gene expression [43–45], several J-Lat clones need to be tested to approximate the polyclonal nature of the latent reservoir *in vivo*, making screenings complicated. To obviate this limitation, we established a polyclonal Jurkat cell model of latency (**Figure 1A**). Briefly, Jurkat cells were infected with a VSV-G-pseudotyped single cycle HIV-1 vector (pNL4.3Δ6-drGFP) [46] (kind gift of Dr. Robert Siliciano, Johns Hopkins University) that expresses a destabilized GFP whose transcription is driven by the viral LTR. This HIV-1 construct has two stop codons in gag-pol and so a separate gag-pol plasmid was provided during transfection of 293T cells to generate the viral supernatants [46]. Infected cells were sorted to select a pure population of GFP+ cells (**Figure 1B**), which were expanded for 5 weeks until latency (GFP- cells) was apparent (**Figure 1C**). Quantitative PCR to detect the GFP gene in the infected cells showed no loss of proviral DNA during passaging (**Figure 1D**), demonstrating actual viral latency.

**Fig 1.**
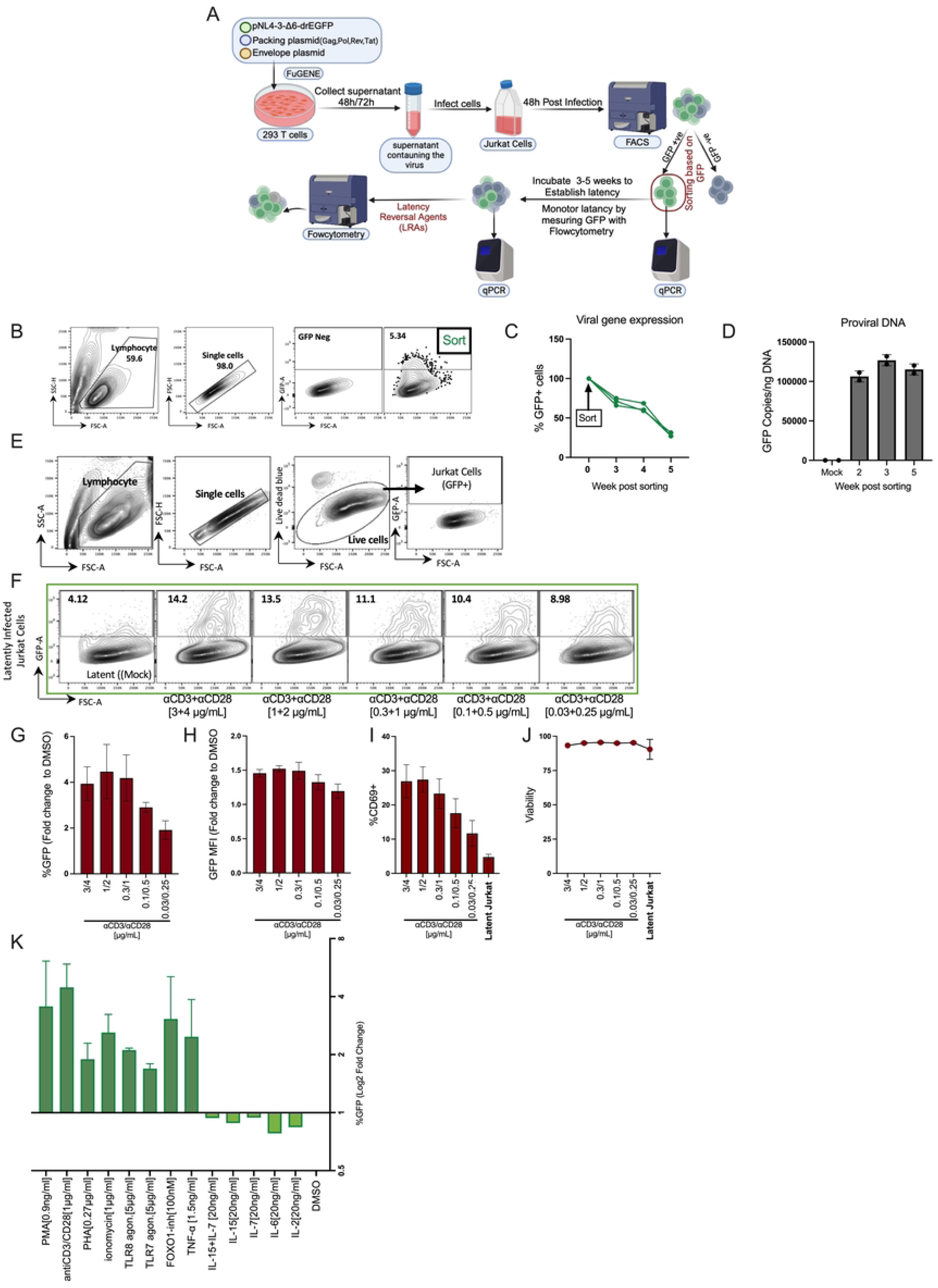
Establishment of a polyclonal Jurkat model of HIV-1 latency. (A) Schematic description of the generation of the latent Jurkat cell model. An HIV-1 reporter virus encoding for GFP under the control of the HIV-1 promoter/enhancer (LTR) was used to differentiate between active and latently infected cells. pNL4-3-Δ6-drEGFP is a ΔEnv, a portion of env was replaced with destabilized GFP (half-life = 4 hours), and ΔNef HIV-1 that has six premature stop codons in gag, vif, vpr, and vpu. The virus only allows for one round of infection. The VSV-G pseudotyped HIV-1 vector was produced by FuGENE transfection into 293T cells. The produced virus was used to infect Jurkat cells, and infection was confirmed by flow cytometry. Forty-eight hours post-infection, GFP+ (infected) cells were sorted by Fluorescence-Activated Cell Sorting (FACS) and maintained until latent infection was established. (B) Flow cytometry gating strategy used for GFP+ cell sorting and monitoring of latency. (C) Flow cytometry analysis for GFP expression at different time points after sorting the infected cells. Three experiments are shown. (D) Quantification by qPCR of proviral DNA copies (using GFP primers) in mock-infected cells and sorted GFP+ cells at different weeks post-sorting. (E) Flow cytometry gating strategy used for detection of GFP+ cells after addition of different LRAs. Uninfected Jurkat cells were used to set the GFP+ gate. (F) Representative flow cytometer plots for anti-CD3/CD28 Ab titration. Latently infected Jurkat cells were activated with five different concentrations of anti-CD3/CD28 Ab or DMSO for 24 hours. The frequency of GFP+ T cells was quantified using flow cytometry; the gating strategy in E was used to detect GFP+ cells. (G) Bar plots showing the percentage of GFP+ cells (fold change to DMSO) after exposure to the indicated concentration of stimuli. (H) Bar graphs showing the GFP MFI fold change to DMSO. (I) Bar graphs showing the % CD69+ cells after anti-CD3/CD28 treatment. Bar blots show average values ± SD, n =6. (J) Viability was measured by flow cytometry using a blue live/dead stain. (K) Summary of latent viral reactivation in Jurkat cells; bar graphs show Log2 fold changes in the percentage of GFP+ cells over DMSO (negative control) for the specified concentration of each LRA. Schematic in (A) was created with BioRender.com.

We then surveyed 14 different LRAs that were previously shown to be active in latently infected primary cells, including cells from PLWH [16, 40, 46–48] (**Table S1**). These LRAs are known to act through either distinct, or partially overlapping mechanisms (**Table S1**) and were therefore suited broadly to probe the pathways for HIV-1 reactivation in our polyclonal latency model. Each LRA was tested at different concentrations to determine the dose-response profile and assess cell toxicity (**Table S2**). Latently infected cells were exposed to the LRAs for 24-48h then analysed by flow cytometry to measure the percentage of GFP+ cells and the GFP mean fluorescent intensity (MFI) as surrogate markers of HIV-1 gene expression. CD69 was used as an early marker of CD4+ T cells activation [49]. Stimulation of the TCR by anti- CD3 and anti-CD28 antibodies showed a dose-dependent increase in both the percentage of GFP+ cells and their MFI relative to untreated cells (**Figures 1E – 1H**). TCR stimulation up- regulated surface expression of CD69, as expected, and did not cause loss of cell viability (**Figure 1I, 1J**). Phorbol 12-myristate 13-acetate (PMA) and Phytohemagglutinin (PHA) also showed a good dose-response in terms of percentage of GFP+ cells, MFI and simultaneous up-regulation of CD69. Ionomycin induced HIV-1 reactivation and CD69 expression in a stepwise fashion above a certain concentration (**Figure S1A-D**).

Toll-like receptor 7 (TLR7) and TLR8 have been reported to reactivate latent HIV-1 in CD4+ T cells by inducing an inflammatory response [48, 50]. We therefore tested if TLR7 and TLR8 agonists promoted reactivation in our model. Treatment of the cells with TLR7 and TLR8 agonists caused a dose-dependent increase in both the percentage of GFP+ cells and MFI but did not significantly affect CD69 expression (**Figure S1C**). FOXO-1 contributes to the maintenance of HIV-1 latency by promoting a quiescent state in infected CD4+ T cells and inhibitors of FOXO-1 can reactivate the latent virus [51, 52]. We therefore tested a well-characterized selective FOXO-1 inhibitor [51, 52] in our cell model and confirmed previous reports by showing that the compound reactivated HIV-1 without affecting cell viability and CD69 expression (**Figure S1E**) [51, 52]. This supports the notion that the HIV-1 and CD69 promoters share some but not all the regulatory pathways [46]. Treatment with TNF-α induced viral reactivation with no CD69 upregulation or loss of cell viability (**Figure S2**) whereas cytokines IL-7, IL-15, IL-6 and IL-2 were inactive in our model **(Figure S2B-D).** The inactivity of IL-7 and IL-15 can be explained by low surface expression of the IL-7 receptor CD127 in Jurkat cells (**Figure S2E**). In summary, 9 known LRAs induced HIV-1 reactivation in our polyclonal model, demonstrating its suitability to test the role of hsp90 (**Figure 1K**).

### Inhibition of hsp90 broadly suppresses HIV-1 reactivation

To test if the LRAs shown in Figure 1K were dependent on hsp90, we employed AUY922, a selective and well characterized hsp90 inhibitor that binds to its N-terminal ATPase pocket and outcompetes ATP [53, 54]. Latently infected cells were treated for 24 hours with a fixed concentration of each LRA in the presence of increasing concentrations of AUY922, or DMSO as control. The LRA concentration chosen had to be within the linear range of the dose-response curve in the absence of cell toxicity. Treated cells were analysed by flow cytometry 24 hours after treatment except for the FOXO-1 inhibitor which was analysed after 48 hours. AUY922 inhibited HIV-1 reactivation mediated by TCR stimulation (**Figure 2A**), PMA (**Figure 2B**), PHA (**Figure 2C**), TNF-α (**Figure 2E**), the FOXO-1 inhibitor (**Figure 2F**) and the TLR7 and TLR8 agonists (**Figure 2G-H**). AUY922 did not appear to be significantly active against ionomycin-induced HIV-1 reactivation even if it antagonized ionomycin-induced upregulation of CD69 surface expression (**Figure 2D**). In general, AUY922 reduced the percentage of GFP+ cells more than the GFP MFI, suggesting that only a proportion of latently infected cells were responding to treatment. This uneven response was reported before in latently infected cells and is presumably related to the stochastic nature of transcription and the chromatin landscape surrounding the provirus [55, 56]. No loss of cell viability was detected in cells treated with both LRAs and AUY922, except in samples treated with the TLR8 agonist and AUY922 at 100nM (**Figure S3**).

**Fig 2.**
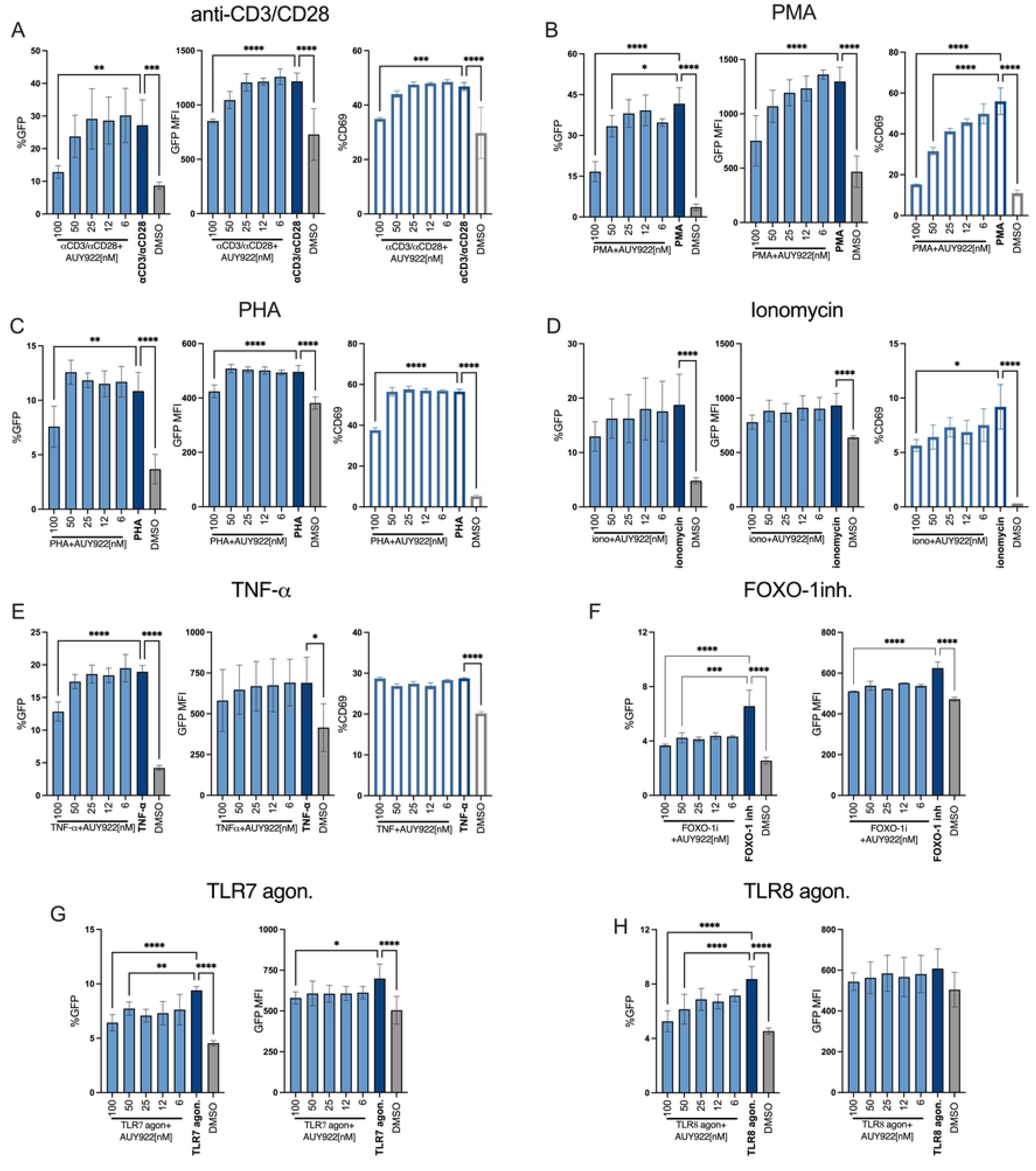
AUY922 represses HIV-1 reactivation in latently infected Jurkat cells. Latently infected Jurkat cells were stimulated with (A) anti-CD3/CD28 Abs [1μg/ml/2μg/ml], (B) PMA [0.9ng/ml], (C) PHA [0.27μg/ml], (D) ionomycin [1 μg/ml], (E) TNF-α [1.5 ng/ml], (F) FOXO-1 inh [200 nM], (G) TLR7 agonist [5μg/ml] (H) TLR8 agonist [5μg/ml], or DMSO to induce HIV-1 reactivation. At the time of stimulation, cells were also treated with the indicated concentrations of AUY922 for 24h and analysed by flow cytometry. In (A-H), bar plots in the left panels show the average percentage ± SD (n=6) of GFP+ cells, middle panels show the average GFP MFI ± SD (n=6), and right panels show the average percentage ± SD (n=3) of CD69+ cells. Statistical significance was calculated using one-way ANOVA with Dunnett’s correction. *=p≤0.05; **=p≤0.01; ***=p≤0.001;****=p<0.0001.

To confirm that these effects were indeed dependent on hsp90 inhibition, we employed the benzoquinone ansamycin 17-allylamino-17-demethoxygeldanamycin (17-AAG or tanespimycin), a well-characterised hsp90 inhibitor that binds to the ATPase pocket of hsp90 but has a different chemical structure from AUY922 (**Figure 3A-B**) [54, 57]. The experimental design with 17-AAG was the same as with AUY22 and selected LRAs were tested. A fixed concentration of LRAs was added to the cells for 24 hours then cells were analysed by flow cytometry to measure the percentage of GFP+ cells, GFP MFI, CD69 and cell viability. 17- AAG antagonised the viral reactivation induced by TCR stimulation, PMA and the TLR7 and TLR8 agonists with no loss of cell viability **(Figure 3C-G)**. Taken together, the results demonstrate that hsp90 broadly regulates HIV-1 reactivation, including that one triggered by TCR stimulation, TLR7 and TLR8 activation and inhibition of FOXO-1, which has not been reported before.

**Fig. 3.**
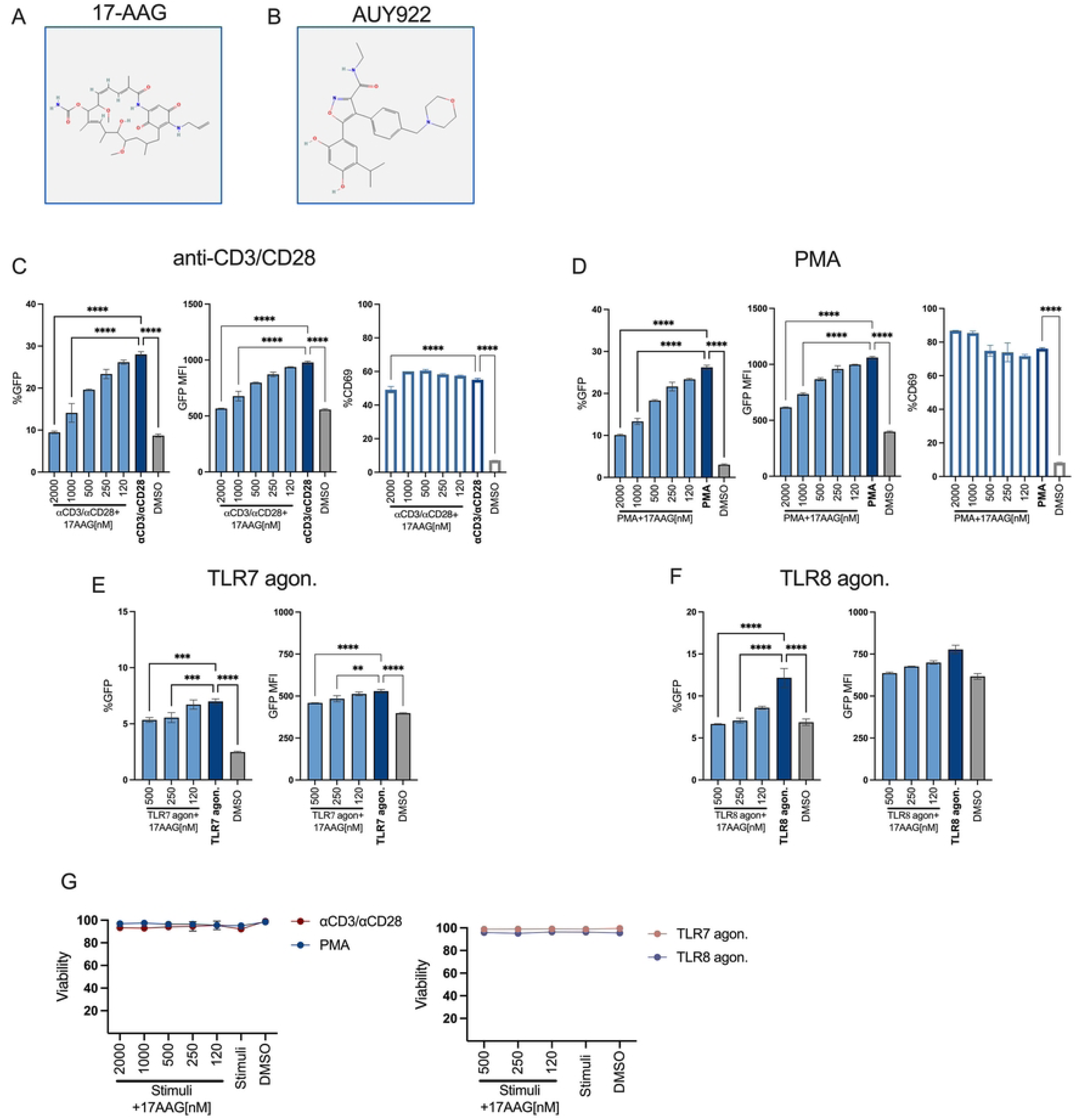
Inhibition of HIV-1 reactivation by 17-AAG in latently infected Jurkat cells. (A) Chemical structure of 17-AAG (PubChem ID: 6505803). (B) Chemical structure of AUY922 (PubChem ID:135539077). (C-F) Latently infected Jurkat cells were stimulated with (C) anti-CD3/CD28 Abs, (D) PMA [0.9ng/ml], (E) TLR7 agonist [5μg/ml] or (F) TLR8 agonist [5μg/ml], or DMSO for 24 hours to induce HIV-1 reactivation. At the time of stimulation, cells were also treated with the indicated concentrations of 17-AAG for 24h and analysed by flow cytometry. In (C-D), left panels show the average percentage ± SD (n=3) of GFP+ cells, middle panels show the average GFP MFI ± SD (n=3) and right panels show the average percentage ± SD (n=3) of CD69+ cells. In (E-F) the left panels show the average percentage ± SD (n=3) of GFP+ cells and the right panels show the average GFP MFI ± SD (n=3). Statistical significance was calculated using one-way ANOVA with Dunnett’s correction. *=p≤0.05; **=p≤0.01; ***=p≤0.001; ****=p<0.0001. (G) Cell viability was monitored by flow cytometry using the live/dead stain.

### AUY922 inhibits the NF-kB, NFAT and AP-1 signal transduction pathways, and this phenotype is recapitulated by targeting TAK1

The broad activity of AUY922 supported the notion that hsp90 regulates multiple signalling cascades important for HIV-1 reactivation. Stimulation of CD4+ T cells by PMA and anti- CD3/CD28 antibodies induce the NF-kB and the AP-1 (c-Jun and c-Fos) signal transduction pathways, which share a key upstream regulator called transforming growth factor-β– activated kinase 1 (TAK1) [58–64] (**Figure 4A**). TAK1 is an hsp90 client protein [65, 66] and has also been reported to bind to the accessory HIV1 protein Vpr, which appears to enhance TAK1 activity [67]. The TCR simultaneously activates PLCγ1 (an hsp90 client protein) and DAG, which, through the PIP(4,5)P_2_, PIP_3_ and Ca^2+^ activates calcineurin and NFAT [68, 69]. NF-kB, AP-1 and NFAT are key regulators of HIV-1 gene expression [19].

**Fig 4.**
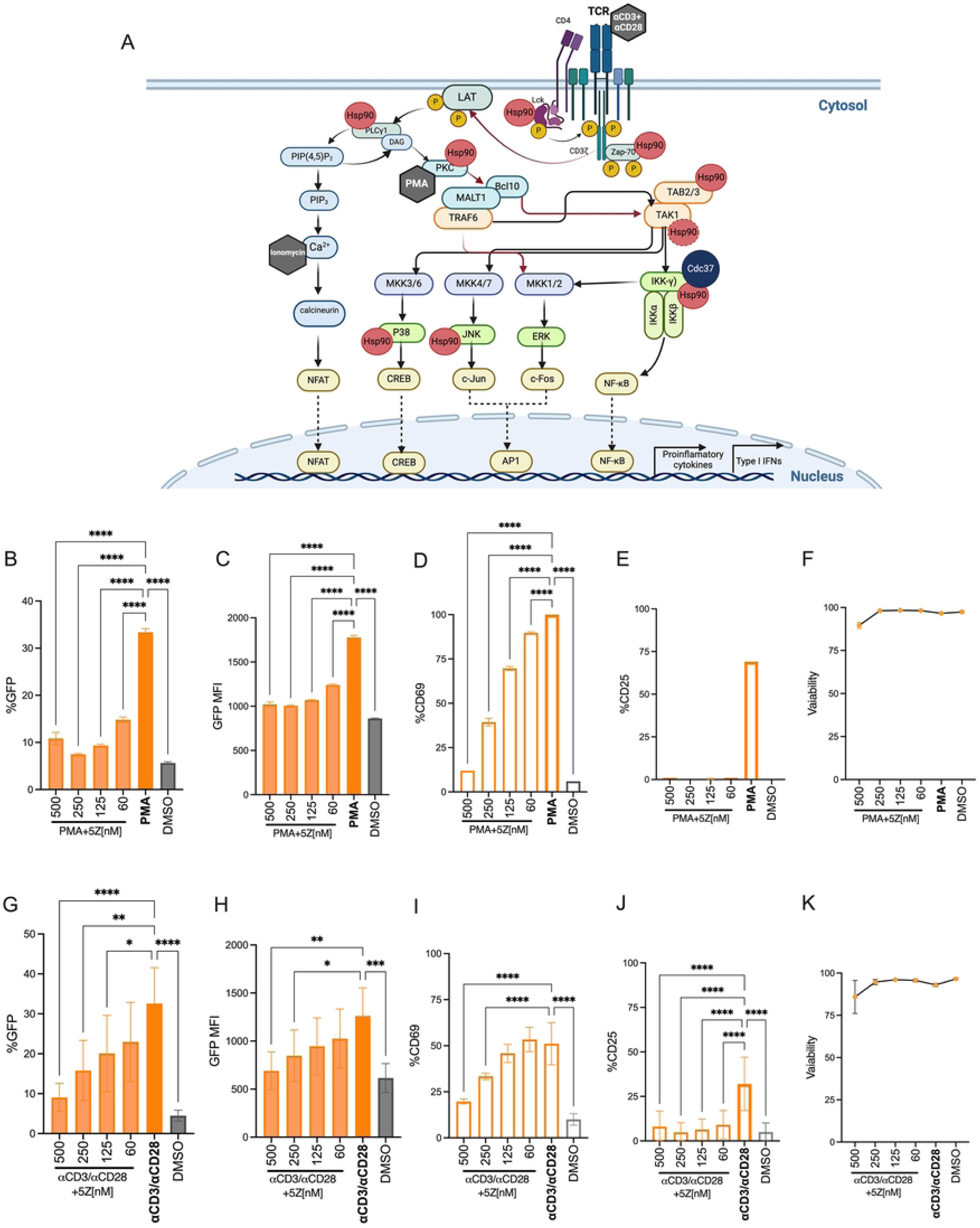
TAK-1 is required for HIV-1 activation. (A) Schematic description of the TCR signal transduction pathways and the factors that are targeted by the LRAs (grey hexagon) or that are known hsp90 client proteins (red circles). (B-F) Latently infected Jurkat cells were stimulated with PMA [0.9ng/ml] for 24 hours in the presence of the indicated concentrations of TAK-1 inhibitor 5Z and analysed by Flow cytometry. (B) Bar graphs show the average percentage ± SD of GFP+ cells, (C) average GFP MFI ± SD, (D) average percentage ± SD of CD69+ cells, (E) average percentage ± SD of CD25+ cells and (F) cell viability. (G-K) Latently infected Jurkat cells were stimulated with anti-CD3/CD28 Abs [1μg/ml/2μg/ml] in the presence of the indicated concentrations of 5Z for 24 hours and analysed by Flow cytometry. (G) Bar graphs showing average percentage ± SD of GFP+ cells, (H) average GFP MFI ± SD, (I) average percentage ± SD of CD69+ cells, (J) average percentage ± SD of CD25+ cells and (K) cell viability. N = 6. Statistical significance was calculated using one-way ANOVA with Dunnett’s correction. *=p≤0.05; **=p≤0.01; ***=p≤0.001; ****=p<0.0001. Schematic in (A) was created with BioRender.com.

We therefore asked if TAK1 regulates HIV-1 latency. To address this question, we stimulated the latent Jurkat cells with a fixed concentration of PMA for 24 hours in the presence of 5*Z*-7- Oxozeaenol (5Z), a selective inhibitor that blocks the ATPase activity of TAK1 [70]. Cells were analysed by flow cytometry to measure the percentage of GFP+ cells, GFP MFI, CD69, CD25 and cell viability. Treatment with PMA triggered viral reactivation, which was repressed by 5Z in a dose dependent way, both in terms of percentage of GFP+ cells and GFP MFI (**Figure 4B-C**). Surface expression of activation markers CD69 and CD25 was also significantly inhibited by 5Z (**Figure 4D-E**) in the absence of detectable cell toxicity (**Figure 4F**). Similar results were obtained following TCR stimulation of the latently infected cells (**Figure 4G-K)**). Hence 5Z mimicked the effect of AUY922 on HIV-1 reactivation.

Next, we sought to test the effect of AUY922 and 5Z on the individual transcription factors NF- kB, AP-1 and NFAT. To this end, we took advantage of the triple parameter reporter (TPR) indicator cells [71], a Jurkat cell line in which the response elements for NF-κB, NFAT and AP- 1 drive the expression of the fluorescent proteins eCFP, eGFP and mCherry, respectively, to simultaneously detect by flow cytometry the activation of these transcription factors after stimulation [71]. Because TPR cells lack the endogenous TCR, here our investigations were limited to stimulation with PMA and ionomycin. Cells were stimulated as before with a fixed concentration of PMA in the presence of AUY922 and analysed by flow cytometry after 24 hours. As expected, PMA triggered robust activation of NF-kB and AP-1, and a more modest activation of NFAT, and AUY922 reduced this effect in a dose dependent way (**Figure 5A-B**). We also tested the combination of PMA + ionomycin for maximal stimulation and found that it enhanced the activation of NFAT and AP-1 above PMA alone, but NF-kB remained unaffected. Inhibition of hsp90 counteracted this effect in a dose-dependent way (**Figure 5C-D**). These results confirmed that hsp90 is critical for the activation of several key transcription factors involved in HIV-1 latency. Of note, the inhibitory effect of AUY922 appeared to be weaker for NFAT relative to NF-kB and AP-1 and to reach a plateau between 25 and 50 nM, suggesting that activation of the NF-kB, AP-1 and NFAT signal transduction pathways does not rely exclusively on hsp90, which may explain the lack of cell toxicity at the tested concentrations.

**Fig 5.**
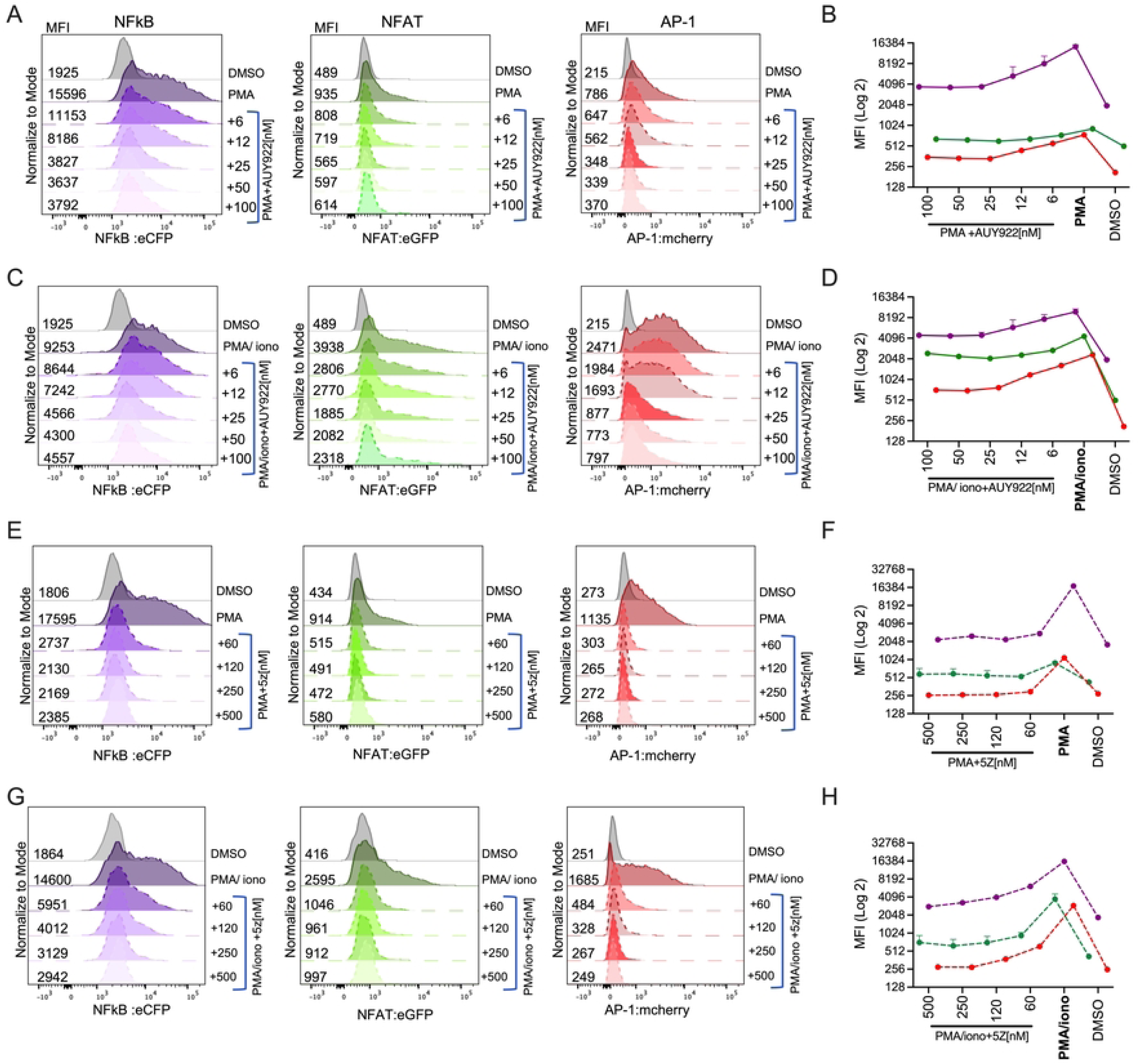
AUY922 and TAK-1 inhibitors suppress NF-kB, AP-1 and NFAT. (A) TPR cells were stimulated with PMA [77 ng/ml] in the presence of the indicated concentrations of AUY922. MFI was measured by Flow cytometry to detect activation of NF-kB (left panel), NFAT (middle panel) and AP-1 (right panel). (B) Graph showing average MFI values ± SD (on a Log2 scale) for each factor, (n= 6). (C) TPR cells were stimulated with PMA [77 ng/ml] + ionomycin [4 μg/ml] in the presence of the indicated concentrations of AUY922 and analysed by Flow cytometry as in (A). (D) Graph showing average MFI values ± SD (on a Log2 scale) for each factor. (E) TPR cells were stimulated with PMA [77 ng/ml] in the presence of the indicated concentrations of 5Z and analysed by Flow cytometry as in (A). (F) Graph showing average MFI values ± SD (on a Log2 scale) for each factor. (G) TPR cells were stimulated with PMA [77 ng/ml] + ionomycin [4 μg/ml] in the presence of the indicated concentrations of 5Z and analysed by Flow cytometry as in (A). (H) Graph showing average MFI values ± SD (on a Log2 scale) for each factor.

We repeated these experiments in the presence of the TAK1 inhibitor 5Z, and the results showed a dose-dependent repression of NF-kB and AP-1 following stimulation with PMA (**Figure 5E-F**). NFAT was less sensitive to 5Z (**Figure 5F**), which mimicked the observations with AUY922 (**Figure 5B**). Stimulation of the cells with PMA + ionomycin enhanced NFAT activity well above PMA alone, and a significant proportion of the signal was reduced by 5Z in a dose-dependent way (**Figure 5G-H**). Taken together, these results indicate that pharmacological inhibition of hsp90 and TAK1 have a similar effect on HIV-1 reactivation, supporting the idea that TAK1 is a key hsp90 chaperone that regulates HIV-1 reactivation.

### Short-term inhibition of hsp90 does not affect the phenotype of primary CD4+ T cells

For clinical applications, agents that suppress HIV-1 reactivation should minimally affect the phenotype and function of the host CD4+ T cells. We therefore investigated if inhibiting hsp90 changed the phenotype and differentiation of primary CD4+ T cells. To this end, CD4+ T cells were isolated by magnetic sorting from peripheral blood mononuclear cells (PBMCs) of healthy volunteers and incubated for 72 hours with IL-2 only (No stim), or anti-CD3/CD28 antibodies plus IL-2 to induce TCR stimulation, or 25 nM AUY922, which was added 48 hours after stimulation. Cells were then analysed by spectral flow cytometry using a 18-colour antibody panel **(Table S3)** [72–78] designed to detect four main CD4+ T cell subsets: naïve (CD3+CD4+, CD45RA+, CCR7+), T central memory (Tcm) (CD3+CD4+, CD45RA-, CCR7+, CD45RO+), T effector memory (Tem) (CD3+CD4+, CD45RA-, CCR7-, CD45RO+), T effector (CD3+CD4+, CD45RA+, CCR7-), and subsets Th17 (CD194+, CD196+), Th1 (CD194- and CD183+), and Th2 (CD194+, CD196-). The activation state of the cells was examined using activation markers CD25, CD69, HLA-DR and CD38 and inhibitory markers PD-1, TIGIT, and Tim-3 **(Table S3)** [72, 79–82]. The gating strategy is described in **(Figure S4)**. To precisely gate the correct population, Fluorescence Minus One (FMO) gating was performed for each of the antibodies on stimulated cells with no AUY922 **(Figure S4).** The parameters set for this FMO staining were subsequently used to gate the samples.

The flow cytometry high dimensional data were analysed by automated t-distributed stochastic neighbour embedding (tSNE) [83]. The tSNE plots showed no appreciable changes in the main cell populations, including samples treated with the hsp90 inhibitor (**Figure 6A and Figure S5**). Expression of the activation markers CD69 and CD25 was higher in stimulated cells and AUY922 reduced CD69 levels in 2 out of 4 donors. No changes were observed for the late stimulation markers HLA-DR and CD38 but treatment with anti-CD3/CD28 Abs downregulated Tim-3 (**Figure 6A, 6B and Figure S5**). We then quantified the flow cytometry results and performed statistical analysis, which confirmed that AUY922 treatment did not significantly change the activation state (**Figure 6B**) or proportion of the different cell populations although we note a trend for fewer CD25+ cells (**Figure 6C**). Induction of CD69 by TCR stimulation was confirmed to be significant, but its downregulation by AUY922 did not show a clear trend. TCR stimulation also resulted in a significant reduction of Tim-3 expression, which was unaffected by AUY922 (**Figure 6C**).

**Fig 6.**
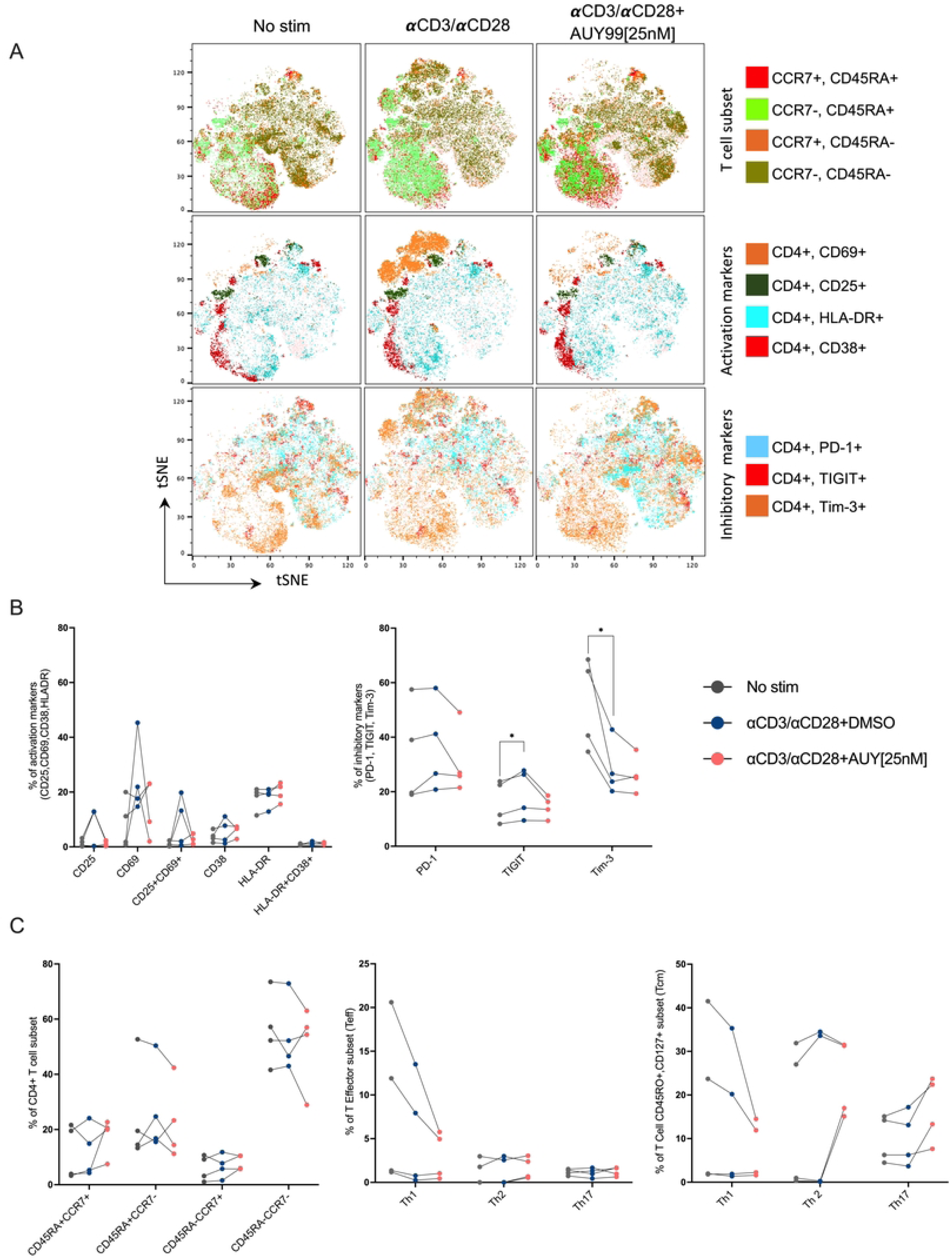
The effect of AUY922 treatment on different CD4+ T cell populations. Primary CD4+ T cells were treated with IL-2 only (No stim) or with anti-CD3/CD28 Abs and IL-2 for 72 hours, and AUY922 [25 nM] or DMSO added at 48 hours post-stimulation. Cells were analysed by spectral flow cytometry using a panel of 18 antibodies 24 hours after the addition of AUY922. (A) Representative tSNE plot for one donor is shown. (B-C) Results from four donors are shown for each CD4+ T cell subset. Subsets were identified according to the combination of surface markers shown in Table S4 and the gating strategy shown in Figure S4. Statistical significance was calculated using two-tailed paired t-Test: *=p≤0.05; **=p≤0.01; ***=p≤0.001; ****=p<0.0001.

### Inhibition of hsp90 suppresses HIV-1 reactivation without affecting the differentiation phenotype of primary CD4+ T cells

These results indicated that inhibition of hsp90 by AUY922 at 25nM did not appreciably alter the overall phenotype and differentiation of uninfected CD4+ T cells (Figure 6). Next, we examined the selectivity of AUY922 in a primary model of HIV-1 latency. To this end, CD4+ T cells were isolated from PBMCs of 5 healthy donors, which were stimulated in vitro by treatment with anti-CD3/CD28 antibodies in the presence of IL-2 for 3 days, after which cells were infected with the single cycle NL4.3Δ6-drGFP virus. Two days later, an aliquot of the cells was analysed by flow cytometry to measure the percentage of GFP+ cells and GFP MFI. Infected cells were maintained in culture with IL-2 for up to 11 days and regularly monitored by flow cytometry to assess the establishment of latency **(Figure 7A-B**). HIV-1 latency manifests as reduction of viral gene expression, best captured in our model by measuring GFP MFI, and also as a complete loss of viral gene expression in individual cells, best captured by measuring the percentage of GFP+ cells. We therefore used both measurements and a combination thereof to assess latency and reactivation in the primary model. In the latency phase, there was a progressive loss of viral gene expression, which was more marked when measured as GFP MFI than percentage of GFP+ cells (**Figure 7C**). Surface expression of CD69+ and CD25+ was also measured to monitor the T cell activation, which showed that infected cells returned to a more resting state over time, except for donor 5 (**Figure 7D**). In parallel, we measured proviral DNA copy number by Alu-LTR qPCR [84] at day 5 and between day 9 and day 11 and found little or no loss of proviral DNA, confirming that genuine latency was established in the primary CD4+ T cell model (**Figure 7E**).

**Fig 7.**
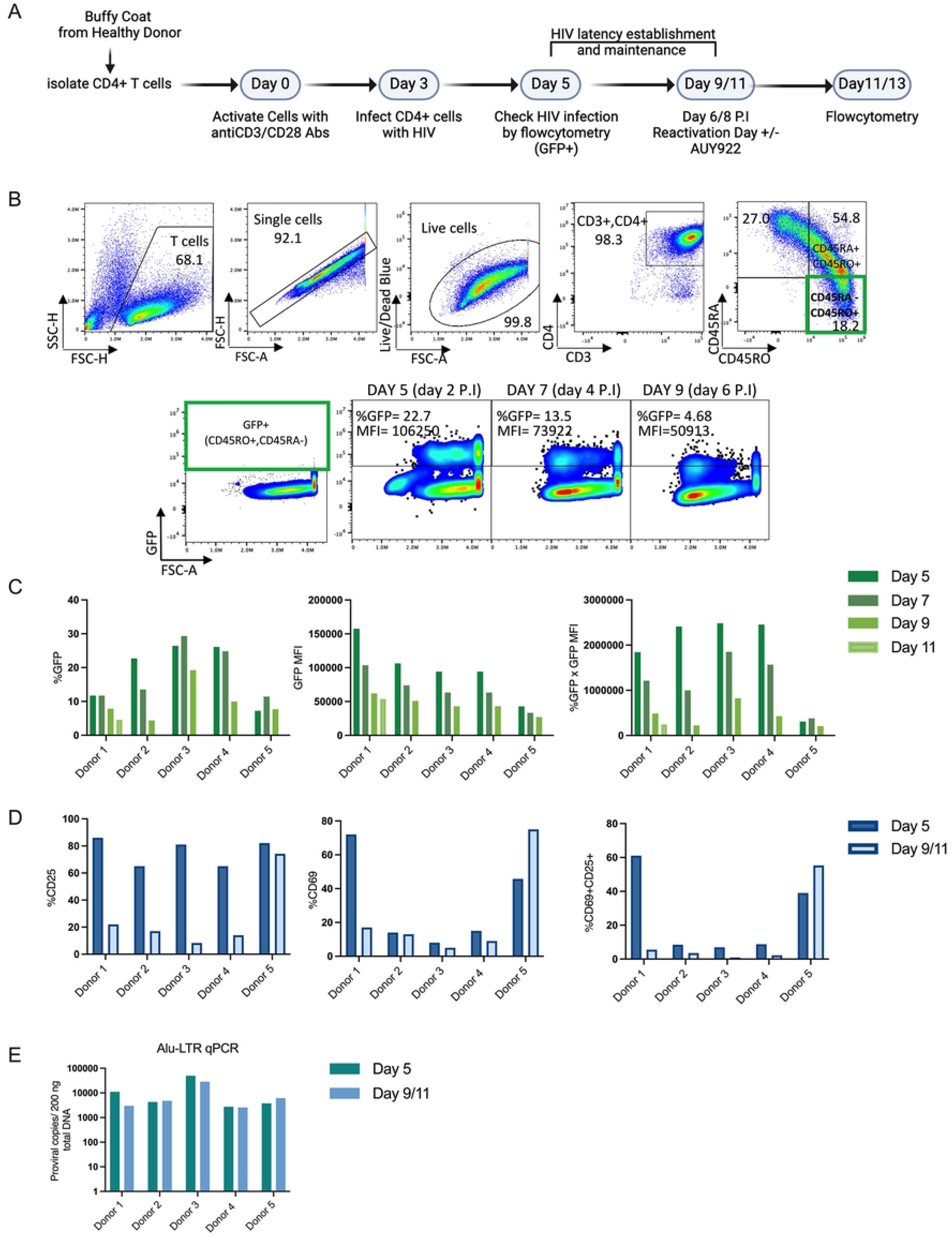
Generation of the ex-vivo latency infected primary CD4+ T cells. (A) Schematic depiction of the experimental set-up to generate latently infected cells ex-vivo. Cells were infected at day 3 post-stimulation with a VSV-G pseudotyped, single cycle HIV-1 reporter virus (pNL4-3-Δ6-drEGFP). (B) Representative flow cytometry plots showing the gating strategy used to detect the GFP+ cells in the CD45RA- CD45RO+ memory population. Cells were analysed by Flow cytometry for GFP expression at regular intervals. (C) Bar graphs showing the GFP MFI (left panel), the percentage of GFP+ cells (middle panel), and combined % GFP x MFI (right panel) measured for each donor on the indicated days. (D) Bar graphs showing the percentage of CD25+ cells (left panel), CD69+ cells (middle panel), and double CD25+CD69+ cells (right panel) for each donor measured by flow cytometry on the indicated days. (E) DNA was extracted from the CD4+ T cells on the indicated days and used to quantify integrated proviral DNA copies by Alu-LTR qPCR. Schematic in (A) was created with BioRender.com.

To induce viral reactivation, cells were exposed to anti-CD3/CD28 antibodies, IL-7/IL-15 or the FOXO-1 inhibitor in the presence of absence of AUY922 (25 or 50nM) or DMSO for 48 hours. Treated cells were analysed by spectral flow cytometry as described in **Figure S4** to measure the percentage of GFP+ cells and its MFI both in the general population and in each CD4+ T cell sub-type, as described in **Figure 7**. In CD45RA- CD45RO+ memory cells, anti- CD3/CD28 antibodies triggered significant reactivation, as measured by the GFP MFI, which was inhibited by AUY922 in a dose-dependent way (**Figure 8A-B**). TCR stimulation did not appear to appreciably increase the percentage of GFP+ cells, nonetheless when both the GFP MFI and the percentage of GFP+ cells were combined, TCR stimulation was found to be effective, and this effect was partially abrogated by AUY922 at 50nM (**Figure 8B**). We then assessed HIV-1 reactivation and the effect of AUY922 in the other CD4+ T cell sub-sets. Within specific CD4+ T cell populations, CD45RA+ CCR7+ naïve cells showed the greatest susceptibility to AUY922, followed by CD45RA- CCR7+ central memory cells [85] (**Figure 8C**), although the latter trend did not reach statistical significance.

**Fig 8.**
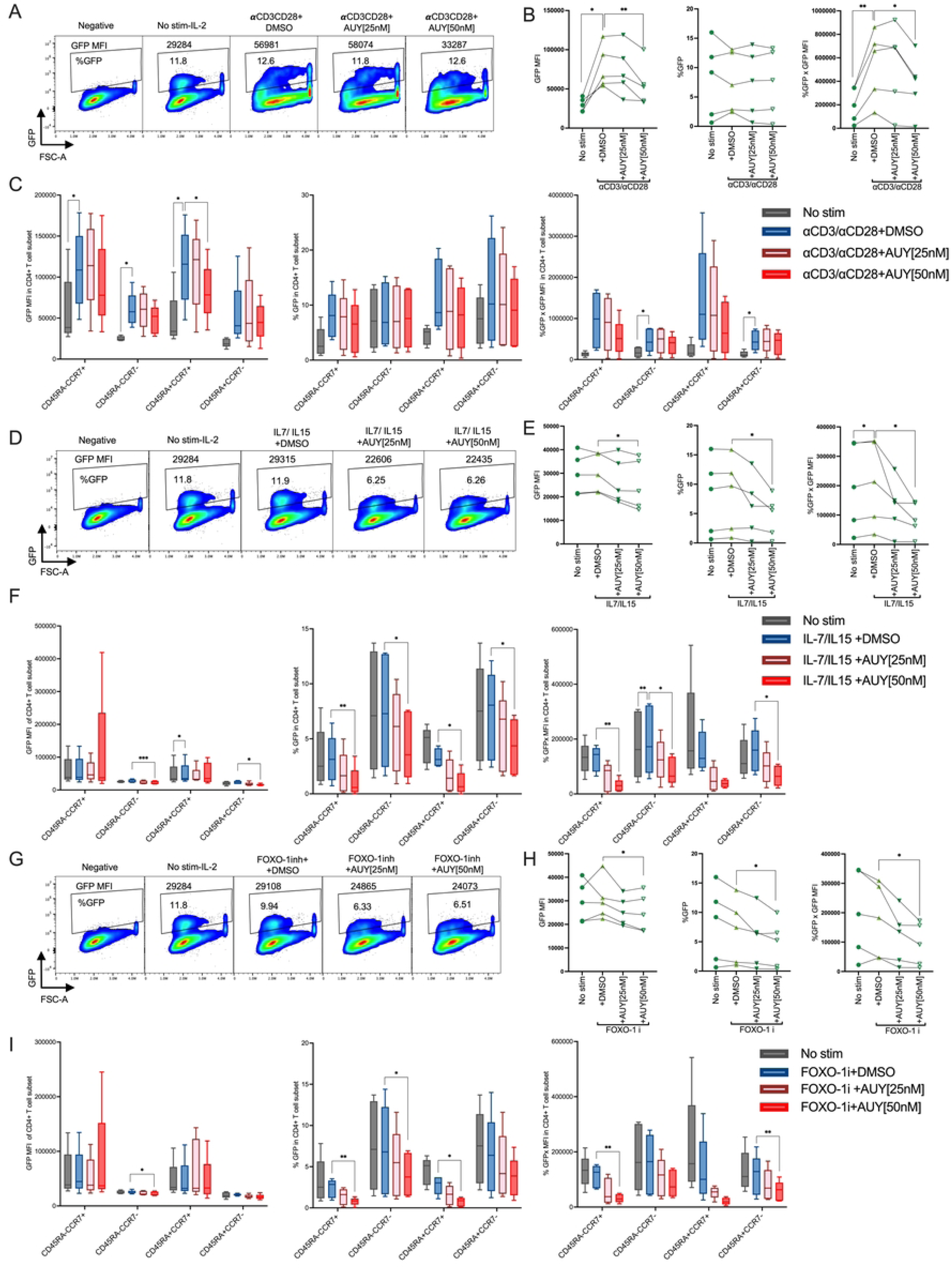
Effect of AUY922 on HIV-1 reactivation induced by different stimuli in primary CD4+ T cell subsets. Latently infected primary CD4+ T cells were generated ex vivo, as described in Figure 7A. Cells were re-stimulated on day 9 or 11 with the indicated LRA with or without AUY922 [25 nM or 50 nM] or DMSO. Cells were analysed by spectral flow cytometry using the same antibody panel used in Figure 6, in addition to GFP. (A) Representative flow plots showing the GFP MFI before (No stim-IL-2) and after re-stimulation with anti-CD3 /CD28 antibodies or AUY922 [25 or 50 nM]. (B) Graphs showing the results for 5 donors: GFP MFI (left panel), percentage of GFP+ cells (middle panel), and combined % GFP x MFI in the CD45RA- CD45RO+ population. (C) Results from five donors are shown for the GFP MFI (right panel), percentage of GFP+ cells (middle panel) and combined % GFP x MFI (right panel) in each CD4+ T cell subset. Bar plots show 1st quartile, 3rd quartile and median for five donors. (D) Representative flow cytometry plots showing the GFP MFI before (No stim- IL-2) and after re-stimulation with IL-7 + IL-15 [20 ng/mL each] with or without AUY922. (E) Graphs showing the results for 5 donors: GFP MFI (left panel), percentage of GFP+ cells (middle panel), and combined % GFP x MFI in the CD45RA-CD45RO+ population. (F) Same as described in (C). (G) Representative flow plots showing the GFP MFI before (No stim-IL-2) and after re-stimulation with the FOXO-1 inh [200 nM] with or without AUY922. (H) Graphs showing the results for 5 donors: GFP MFI (left panel), percentage of GFP+ cells (middle panel), and combined % GFP x MFI in the CD45R-CD45RO+ population. (I) Same as described in (C). Statistical significance was calculated using a two-tailed paired t-Test comparing anti-CD3CD28 versus no stim-IL-2, and anti-CD3CD28 versus 50 nM AUY922, *=p≤0.05; **=p≤0.01; ***=p≤0.001; ****=p<0.0001.

Cells treated with IL-7 and IL-15 did not show appreciable reactivation, however inhibition of hsp90 significantly reduced viral gene expression below the baseline (unstimulated cells) in CD45RA- CD45RO+ memory cells and this effect was significant for GFP MFI, percentage of GFP+ cells and the combination of GFP MFI and % GFP+ cells (**Figure 8D-E**). Within specific CD4+ T cell subsets, the greatest susceptibility to AUY922 was detected in CD45RA+ CCR7+ cells, followed by CD45RA- CCR7+ cells, and CD45RA- CCR7- effector memory cells (**Figure 8F**). Treatment with the FOXO-1 inhibitor failed to induce viral reactivation but inhibition of hsp90 significantly reduced viral reactivation below the baseline of unstimulated CD45RA- CD45RO+ cells (**Figure 8G-H**). In the FOXO-1 inhibitor-treated cells, we detected the strongest response to AUY922 in the CD45RA- CCR7+ and CD45RA+ CCR7- populations, followed by the CD45RA+ CCR7+ population (**Figure 8I**).

It is not clear why IL-7 + IL-15 and the FOXO-1 inhibitor did not trigger appreciable viral reactivation however we note that the conditions of the experiments on Jurkat cells and on primary CD4+ T cells were different, mainly because the primary cells had been pre-stimulated with anti-CD3/CD28 antibodies to make them permissive to HIV-1 infection 10 days before re- stimulation, which might have affected their response to the LRAs themselves. Furthermore, the threshold for the response to LRAs might be different between Jurkat and primary cells. Nonetheless, the results confirmed that targeting hsp90 reduced TCR-induced reactivation and inhibited baseline viral gene expression in latently infected cells even in the presence of IL-7 and IL-15 or the FOXO-1 inhibitor.

We also examined if inhibition of hsp90 at levels sufficient to reduce viral gene expression affected the differentiation and phenotype of CD4+ T cells. In cells stimulated twice by anti- CD3/CD28 antibodies, we found a significant upregulation of activation markers CD69, CD25, CD38 and CD71 (the transferrin receptor) [86] with simultaneous upregulation of inhibitory markers PD-1, TIGIT and Tim-3 (the latter showed a trend but did not reach statistical significance), which suggested a degree of exhaustion after two rounds of stimulation in a relatively short time interval [80] (**Figure 9A**). Treatment with AUY922 did not appreciably affect expression of these markers, except for a reduction in TIGIT surface expression (**Figure 9A**). We also found a significant reduction in the proportion of markers of Th1 cells against a significant increase in the proportion of markers of Th2 cells but treatment with AUY922 did not change this (**Figure 9A**). Treatment with IL-7/IL-15 significantly upregulated HLA-DR and CD71, confirming activity of the cytokines, and AUY922 significantly reduced expression of CD71 (**Figure 9B**). The FOXO-1 inhibitor significantly upregulated CD25, downregulated CD69 and showed a trend for CD71 upregulation (**Figure 9C**). These results indicated that IL- 7/IL-15 had biological activity on the CD4+ T cells, even if they did not stimulate HIV-1 reactivation. Therefore, in our experimental conditions, inhibition of hsp90 did reduce HIV-1 gene expression without any apparent significant effect on the differentiation and phenotypic markers of CD4+ T cells.

**Fig 9.**
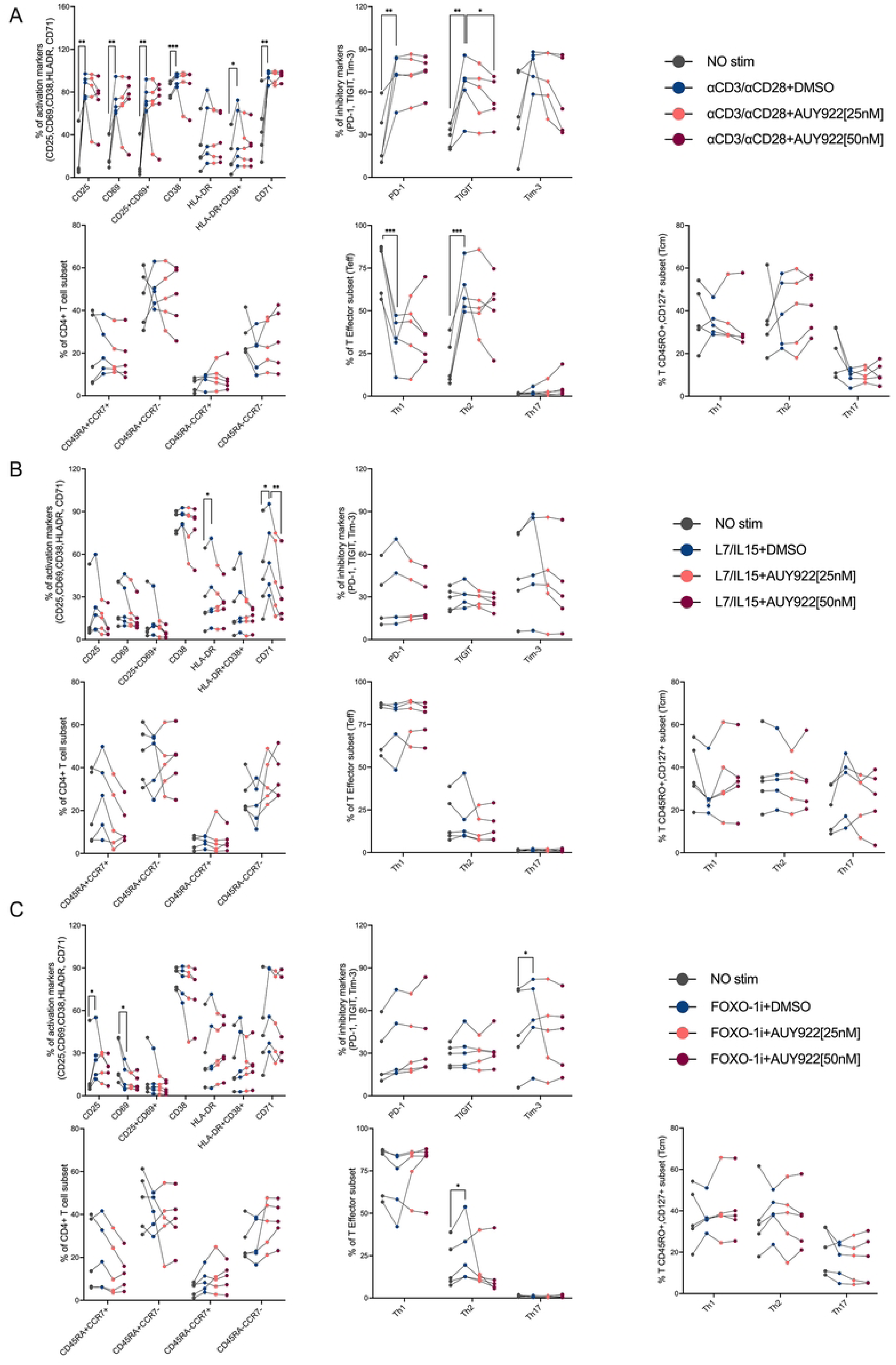
AUY922 does not significantly change the CD4+ T cell phenotype or activation state. Latently infected primary CD4+ T cells were re-stimulated on day 9 or 11 with the indicated LRA with or without AUY922 [25 nM or 50 nM] or DMSO. Cells were analysed by spectral flow cytometry using the same antibody panel used in Figure 6 with the addition of CD71. (A) panels show, in this order from left to right, top to bottom: T cell activation markers, inhibitory markers, CD4 T cell subsets, effector subsets and memory subsets after re-stimulation with anti-CD3 [1 μg/ml] and anti-CD28 [2 μg/ml] antibodies plus AUY922 [25 or 50 nM] or DMSO. (B) Same measurements for cells re-stimulated with IL-7/IL-15 [20ng/mL]. (C) Same measurements for cells restimulated with FOXO-1 inh. [200nM]. Results were obtained from five different donors. Statistical significance was calculated using a two-tailed paired t-Test comparing, within each cell sub-type, anti-CD3CD28 versus no stim (IL-2), anti-CD3CD28 versus 50 nM AUY922, *=p≤0.05; **=p≤0.01; ***=p≤0.001; ****=p<0.0001.

Lastly, we investigated if treatment with AUY922 affected cytokines production from activated CD4+ T cells. To this end, we collected the supernatant from the cell cultures of the same 5 donors described in Figure 8B. Supernatants were collected 48h post-restimulation with anti- CD3/CD28 Abs to measure the concentration of 5 cytokines that define specific subsets of effector CD4+ T cells: IFNγ and TNF-α (Th1), IL-4 and IL-10 (Th2) and IL-17A (Th17) [87]. TCR stimulation significantly increased production of all tested cytokines, however AUY922 significantly but modestly inhibited only IL-4 and IL-10 production (**Figure S6**). There was a downward trend for the other cytokines in the AUY922-treated samples compared to untreated, but it did not reach statistical significance. These results confirmed that inhibition of hsp90 did not broadly perturb CD4+ T cell function, although it might modestly reduce production of Th2 and Th1 cytokines.

## Discussion

HIV-1 latency is regulated at multiple levels, which include the epigenetic features of chromatin at the proviral integration site, the availability of specific host cell factors that activate viral transcription, the post-translational regulation of such host factors and the intactness of the provirus itself [18, 19]. The mechanisms that influence HIV-1 latency, or its reactivation, are intimately linked to the activation and differentiation of the infected CD4+ T cells [19]. Therefore, strategies to either enhance HIV-1 reactivation from latency or induce a deeper state of latency need to address two problems: the multi-factorial nature of latency and simultaneously interfering as little as possible with the function and differentiation of the infected cells.

Here, we have confirmed that inhibiting hsp90 antagonises HIV-1 reactivation triggered by LRAs such as PMA, PHA and TNF-α [13] and showed, for the first time to our knowledge, that hsp90 is also required for HIV-1 reactivation induced by TCR, TLR7 and TLR8 stimulation and FOXO-1 inhibition. No significant loss of cell viability was detected in our experimental conditions and similar results were obtained using AUY922 or 17-AAG, two structurally different hsp90 inhibitors that bind to the same ATPase pocket in the N-terminal region of the chaperone, supporting the specificity of the effect. AUY922 was chosen for our experiments because it is a well-characterized drug that has been used in phase II and III clinical trials [88].

The repressive effect of AUY922 was broad and extended to many pathways that are physiologically involved in HIV-1 reactivation [89, 90]. This can be explained by the numerous client proteins that depend on hsp90 for correct folding and assembly, which are also key components of the signalling pathways critical for virus reactivation [91]. We and others have previously shown that hsp90 is part of the IKK complex and promotes its activity, and that it stabilizes P-TEFb [13, 30]. Here, we have extended these observations and shown that hsp90 is critical for activation of the NF-kB, and AP-1 signal transduction pathways and to a lesser extent the NFAT pathway, supporting the notion that the chaperone controls multiple key events in viral reactivation. Notably, we found that inhibition of TAK1, an hsp90 client protein, by the selective antagonist 5Z phenocopied AUY922, suggesting that at least part of its anti- reactivation effect is mediated by TAK1.

Hence short-term inhibition of hsp90 appears to address, albeit not fully, the first problem related to the multifactorial nature of HIV-1 latency, and in this respect hsp90 may be considered a master regulator. But does inhibition of hsp90 addresses the second problem related to the effect of latency potentiating agents on the physiology of the infected CD4+ T cells?

To answer this, we have initially examined the effect of AUY922 on the phenotypic profile of uninfected CD4+ T cells. A well-established method to study the profile of T cells and their differentiation is multiparameter flow cytometry, which measures the combinatorial expression of chemokine receptors and other differentiation markers on the cell surface [72]. Surface expression of these markers is tightly regulated at the transcriptional and post-transcriptional levels and reflects the overall physiological state of the cell [92, 93]. We applied a carefully selected panel of 18 markers that, when suitably combined, define several subsets of CD4+ T cells, including naïve, effector, central memory, effector memory and subtypes Th1, Th2 and Th17. We have also employed surface markers of activation, whose expression changed depending on the applied LRA in agreement with its known physiological effect. The results of these experiments revealed no significant change in any of the cell phenotypes, suggesting that the degree of hsp90 inhibition applied to inhibit HIV-1 reactivation was well tolerated.

Next, we assessed the impact of AUY922 on infected CD4+ T cells. We generated ex-vivo primary latently infected cells, which required a first round of T-cell stimulation to make cells permissive to HIV-1 infection, passaging the cells to allow them to return to a more quiescent state and establish viral latency, and a second round of stimulation to trigger viral reactivation in the presence of AUY922. We were able to establish viral latency in this model, although there was donor-to-donor variability in the magnitude of reactivation. AUY922 antagonized viral reactivation and reduced the level of viral gene expression as measured by the GFP MFI, although it did not seem to fully block it, which would have resulted in a lower percentage of GFP+ cells. This differed from Jurkat cells, in which AUY922 reduced both GFP MFI and percentage of GFP+ cells, suggesting greater potency of the drug. This discrepancy between primary CD4+ T cells and Jurkat cells may be due to the well-known heightened sensitivity of cancer cells to hsp90 inhibitors [94, 95].

The combination of IL-7 and IL-15 did not reactivate HIV-1 in our primary model or in Jurkat cells. IL-7 renders resting CD4+ T cells more permissive to HIV-1 infection [96] through the Jak/STAT5 pathway [97] and by partially activating the cells [98]. IL-7 has also been reported to activate latent HIV-1 ex-vivo in human CD4+ T cells from humanized mice [99], and in cells from patients at the same concentrations we tested, however this effect was reported to be variable from donor to donor, proviral-strain specific and was seen after several days of exposure to the cytokine [41, 100]. Furthermore, significant IL-7 induced reactivation is not universally observed [101, 102]. We detected upregulation of HLA-DR, as previously described [100], and CD71 upon treatment with IL-7/IL15, but we did not detect viral reactivation. We surmise that the difference with some earlier studies may be due to the shorter duration of treatment in our experiments relative to other studies [41, 100]. Notably, AUY922 suppressed viral gene expression below the baseline even in the presence of IL-7/IL- 15, confirming its activity on the HIV-1 promoter [13].

Treatment of the cells with the FOXO-1 inhibitor upregulated surface expression of CD71, in agreement with previous reports [52] but induced weak reactivation in two out of five donors. This result contrasted with the reproducible reactivation that we and others found in Jurkat cells [51, 52] and we speculate that this difference in the response is due to the short time of exposure to the FOXO-1 inhibitor. Regrettably, our primary latency model did not allow to extend the duration of the experiment beyond 12-13 days because of a notable drop in cell viability after that time point. Nonetheless, even in the presence of the FOXO-1 inhibitor, AUY922 reduced viral reactivation below the baseline of untreated cells.

Different cell subsets showed different responses to the LRAs and AUY922. Overall, the strongest viral reactivation after TCR stimulation and the greatest susceptibility to AUY922 was detected in CD45RA+ CCR7+ cells, which is largely made of naïve cells but can also contain a small proportion of T memory stem cells (Tscm) [103] followed by CD45RA- CCR7+ central memory cells, and CD45RA- CCR7- effector memory cells, which agrees with a previous study [104]. Certain CD4+ T cell subsets have been shown to be more susceptible to specific LRAs but the reasons for this behaviour are not clear [105–108]. Our results extend these observations to include CD4+ T cell subtype-specific responses to hsp90 inhibition and it will be interesting to investigate the mechanistic reason for this phenotype and how different LRAs and hsp90 intersect in the different CD4+ T cell subtypes.

Notably, treatment with AUY922 did not significantly change the proportion of each CD4+ T cell subtype in the population and had a modest effect on Th2 and Th1 cytokine production, which demonstrated that inhibiting hsp90 does reduce HIV-1 reactivation without dramatically altering CD4+ T cell differentiation or activation state, at least in the experimental conditions tested. The weak inhibition of Th1 cytokine production may be advantageous in PLWH, who often have heightened general inflammation. We note that TCR stimulation enhanced surface expression of activation and exhaustion markers and reduced the relative proportion of the markers for Th1 cells, indicating that our readout was able to detect phenotypic changes in the samples. Considering that hsp90 participates in a wide variety of cellular pathways, these results may seem surprising. Selectivity is usually a function of the drug concentration and time of exposure, which in our case was limited to 48 hours. It is possible that a longer incubation time in the presence of the drug may affect the phenotype of CD4+ T cells, and it should be possible to evaluate this aspect in patients who are being treated with AUY922 or other hsp90 inhibitors in clinical trials [109].

Hsp90 inhibitors are being evaluated in clinical trials for the treatment of both solid and haematological malignancies, at concentrations that are significantly higher than those that are sufficient to repress HIV-1 gene expression [88]. Although so far hsp90 inhibitors have not shown good efficacy above baseline for the treatment of solid cancers [109, 110], their pharmacological and toxicity profiles are well-known. It would therefore be conceivable to assess hsp90 inhibitors in the context of analytical treatment interruption (ATI) studies to test if virological rebound is delayed with the view of using the inhibitors in conjunction with other eradication strategies, especially if the long-lasting repression of virological rebound reported in humanized mice are reproduced in patients [28]. An interesting question to address in future studies will be whether inhibition of hsp90 synergises with the tat inhibitor TCA [24, 25] or with AhR agonists [21] to stably repress HIV-1 reactivation.

Lastly, hsp90 is required for the replication of many RNA and DNA viruses [111], including early gene expression of human cytomegalovirus (HCMV) [112] and it could therefore be a broad antiviral target for the treatment of co-infections, often seen in PLWH [113].

Our study has some limitations. The Jurkat and primary cell models of latency may not fully recapitulate the complexity of the in vivo latent reservoir and the viral construct used does not express accessory proteins Vpr and Vpu, which may have a role in latency and viral gene expression [114, 115]. Although we observed clear trends in the primary cell model, not every result reached statistical significance due to donor-to-donor variability in the degree of viral reactivation and this issue will need to be carefully considered in future pre-clinical studies.

## Materials and Methods

### Ethics statement

Blood samples were obtained from healthy volunteers after written informed consent, or from the National Health Service Blood and Transplant (NHS-BT) according to the approved protocol of the University College London Research Ethics Committee reference REC 3138/001 and NHS-BT reference R140.

### Chemical compounds

NVP-AUY922 was purchased from LKT Laboratories; 17-allylamino-17- demethoxygeldanamycin (17-AAG or tanespimycin) from Merck Life Science UK Ltd. Phorbol 12-myristate 13-acetate (PMA), 5*Z*-7-Oxozeaenol (5Z) were obtained from Cayman Chemical; TNF-a was purchased from Life Technologies Limited. Imiquimod (TLR7 agonist) was purchased from SAlfaAesar, CL075(TLR8 agonist) from Sigma-Aldrich (MREK). FOXO-1 inh (AS1842856) was purchased from MedChemExpress LLC, Phytohemagglutinin (PHA) and Ionomycin were obtained from Thermo Fisher Scientific. Compounds were dissolved in DMSO to obtain 1000x stock solutions (v/v) and stored in aliquots at −20 °C in the dark. Purified anti-human CD3 antibody (OKT) and anti-human CD28 antibody (CD28.2) were purchased from Biolegend; Recombinant human IL-7, rhIL-2, IL-6 and IL-15 were obtained from Peprotech. The concentrations used are shown in Table S2 for each compound.

### Cells and tissue culture

The Jurkat cell line E6-1 was obtained from the American Type Culture Collection (ATCC) and cultured in RPMI media (Gibco) supplemented with 10% fetal bovine serum, 100 U/mL penicillin/streptomycin, and maintained at 37°C in a 5% CO₂ incubator. 293T cells were obtained from the ATCC and maintained in DMEM media (Gibco) supplemented with 10% fetal bovine serum, 100 U/mL penicillin/streptomycin. Peripheral blood mononuclear cells (PBMCs) were isolated by Ficoll-Paque PLUS density gradient centrifugation. Human CD4+ T lymphocytes were isolated from PBMCs using the MojoSort Human CD4+ T Cell Isolation Kit (BioLegend, 480010) following the manufacturer’s instructions. Isolated CD4+ T cells were cultured in X-VIVOTM 15 Serum-free Hematopoietic Cell Medium (Lonza), supplemented with 5% fetal bovine serum (FBS; Labtech) and 100 U/mL penicillin/streptomycin. Triple parameter reporter (TPR) Jurkat-derived cells were kindly provided by Prof. Peter Steinberger (Medical University of Vienna) and cultured in RPMI as above.

### Virus production

The single-cycle HIV-1 vector (Δenv) was produced by transfection of 293T cells using FuGENE Transfection Reagent as previously described [22, 45]. The transfection mixture included the following plasmids: pNL4-3-Δ6-drEGFP, pCMV-VSV-G envelope, and pCMV-Gag/Pol packaging vectors. Supernatants containing the virus were collected 48 and 72 hours post-transfection, filtered through a 0.45 µm filter, and concentrated by ultracentrifugation at 25,000 rpm for 2 hours at 4°C through a 25% sucrose cushion [116]. The viral pellet was resuspended in RPMI medium and stored at -80°C. The virus stock titer was determined by infecting Jurkat cells with serial dilutions of viral-containing supernatants and detection of GFP expression by flow cytometry two days after cell transduction.

### Generation of latently infected Jurkat cells

Jurkat cells were infected with NL4.3Δ6-drGFP viral stock at an MOI of 0.2. Forty-eight hours post-infection, GFP+ (infected) cells were sorted using BD FACSAria II and the GFP+ population was maintained in culture and regularly monitored by flow cytometry for GFP expression until latent infection was established (2-3 weeks). For stimulation of latently infected Jurkat cells, 1 × 10⁵ cells were treated with the indicated concentrations of LRAs (**S2 Table**) for 24 hours, except for the FOXO-1 inhibitor, which was added for 48 hours. Cells were then analysed by flow cytometry to measure the percentage of GFP+ cells and the GFP MFI. AUY922, 17-AAG and 5Z-7-Oxozeaenol (TAK-1 inhibitor) were dissolved in DMSO and used at serial dilutions as indicated in the Figures. To assess the toxicity of the inhibitors, cells were stained with LIVE/DEAD Fixable blue dead cell stain (Thermo Fisher scientific) before analysis. For all Jurkat cells experiments, fluorescence was measured on a BD LSR Fortessa and analysed using the FlowJo software 10.8.1.

### Infection of primary CD4+ T cells and generation of latently infected cells

CD4+ T cells were activated by anti-CD3/CD28 monoclonal antibodies. Tissue culture plates were precoated with anti-CD3 (OKT) Ab at 1 µg/mL then cells were added to the plate together with soluble anti-CD28 Ab (CD28.2) at 2 µg/mL and 100 U/mL IL-2 in X-VIVO media supplemented with 5% FBS and 100 U/mL penicillin/streptomycin. On day 3, the activated CD4+ T cells were infected with the NL4.3Δ6-drGFP virus at an MOI of 2 in the presence of 4 µg/mL polybrene and 100 U/mL IL-2. The cells were incubated overnight on a shaker in the incubator. Cells were removed from the shaker, incubated for 24 hours then cells were stained for CD3, CD4+, CD45RA, CD45RO, CD25, and CD69 and analysed by flow cytometry. To reactivate latently infected CD4+ T cells, 1 × 10⁵ cells were cultured in 96-well plates in 100uL media and treated with different LRAs, including anti-CD3/CD28 (1 µg/mL/2 µg/mL), IL-7 + IL- 15 (20 ng/mL of each), AS1842856 (200 nM), TLR7 agonist (5 μg/mL), and TLR8 agonist (5 μg/mL). Cells were treated with each LRA in the presence of either DMSO, AUY922 (25 nM), or AUY922 (50 nM) and incubated for 48 hours before staining and analysis by flow cytometry. Additionally, the supernatant of these cells was collected for cytokine analysis.

### Cell Surface Staining for Flow Cytometry

For surface marker staining, the appropriate human-specific mAbs were chosen (see Table below). For Jurkat cells and TPR cells, fluorescence was measured using a BD LSR Fortessa

,BD FACS Diva9 software, while for infected CD4+ T cells, fluorescence was measured using a Cytek Aurora. Data were analysed using FlowJo version 10.8.1. The gating strategy used to identify T cells subsets is shown in Fig. S4

| Fluorochrome | Specificity (Marker) | Supplier (catalogue number) |
| --- | --- | --- |
| BV510 | TIGIT | BD (747842) |
| BV711 | Tim-3 | BD (747959) |
| BV786 | CD197(CCR7) | BD (566758) |
| BV421 | CD196(CCR6) | BD (562515) |
| BV605 | CD71 | BD (745096) |
| APC/fire 810 | CD25 | BioLegend (356150) |
| APC | CD69 | Biolegend (310910) |
| Alexa fluor 700 | CD 127 | BioLegend (351344) |
| APC/Cyanine 7 | CD4+5RO | BioLegend (304228) |
| BUV805 | HLA-DR | BD (748338) |
| UV (450) live dead blue | Live/Dead | ThermoFisher (L34961) |
| BUV 395 | CD4+5RA | BD (740315) |
| BUV737 | CD183 | BD (741866) |
| Spark NIR 685 | CD38 | BioLegend (303552) |
| PE-Dazzle | PD-1 | BioLegend (367434) |
| PE cy7 | CD4+ | Biolegend (317414) |
| PE/fire 810 | CD194(CCR4) | BioLegend (359433) |
| PE | CTLA4(CD152) | BioLegend (369604) |
| GFP | GFP reporter |  |
| Percp cy5.5 | CD3 | Biolegend (317336) |

### Quantification of HIV-1 integration

To quantify the integration of HIV-1 in CD4+ T cells, nested Alu-LTR quantitative PCR was performed as previously described [112]. Briefly, DNA was isolated from infected CD4+ T cells 48 hours post-infection, 10 days post-infection, and from a negative sample (uninfected) using the Qiagen Blood Mini Kit. Integrated DNA was pre-amplified using 50 nM Alu forward primer, 150 nM HIV-1 LTR reverse primer, 25 μL PCR Master Mix (2X) (ThermoFisher), and 200 ng DNA in 50 μL reactions. Cycling conditions were: 95°C for 8 min x 1 cycle, followed by 18 cycles of 95°C for 1 min, 52°C for 1 min, and 72°C for 3 min. A second round real-time TaqMan quantitative PCR was performed using the pre-amplified DNA. These samples were run alongside a standard curve of known dilutions of infected Jurkat cells containing integrated HIV-1 DNA. Reactions contained 0.5 μM Alu forward primer, 0.5 μM Alu-LTR2 reverse primer, 0.15 μM probe, 10 μL 2x TaqMan Gene Expression Master Mix, and 2 μL of 1:20 diluted pre-amplified DNA. Cycling conditions were: 50°C for 2 min, 95°C for 15 min x 1 cycle, followed by 45 cycles of 95°C for 15 s, and 60°C for 1 min. Reactions were performed using a QuantStudio real-time PCR system. Below is the list of primers used: ALU-FW: AAC TAG GGA ACC CAC TGC TTA AG LTR1-RV: TGC TGG GAT TAC AGG CGT GAG LTR2-RV: TGC TAG AGA TTT TCC ACA CTG ACT ALU- Probe: FAM—TAG TGT GTG CCC GTC TGT TGT GTG AC—TAMRA

### qPCR for proviral quantification

Total DNA from 1.5 × 10^6^ infected WT or N74D Jurkat cells was extracted at each week after sorting using DNeasy blood & tissue kit (Qiagen). DNA concentration and purity were measured by Nonodrop and each DNA sample was normalized to 100ng/μl. The real-time qPCR reaction was performed using a Real-Time PCR machine (Applied Biosystems) in a final volume of 20 μl containing 1x Power-Up SYBR green master mix (Applied Biosystems), 200ng of DNA in 2 μl and 0.2 μM of each primer: forward GFP primer AAGCTGACCCTGAAGTTCATCTGC and reverse GFP primer CTTGTAGTTGCCGTGGTCCTTGAA. Cycling parameters were 95^0^C for 2 min followed by 95^0^C for 1 min, 55^0^C for 1 min and 68^0^C for 1 min repeated for 40 cycles. Quantification was done using a standard curve that was generated by serial dilutions of the Δ6-drEGFP plasmid.

### Cytokine analysis

Supernatants from CD4+ T cells isolated from five different donors were collected on the day of cells analysis. These supernatants were then analysed for IFN-γ, IL-4, IL-17A, IL-10, and TNF-α levels using the LEGENDplex™ Human Essential Immune Response Panel Mix and Match (Cat #740932, BioLegend) following the manufacturer’s protocol. The measurements were taken using a BD LSRFortessa flow cytometer, and the data were analysed with the LEGENDplex™ Data Analysis Software.

### Statistical analysis

Means ± SE or SD, n, and the statistical test used are shown in the Figure legends. Statistical analyses were conducted using GraphPad Prism software. The levels of statistical significance are indicated as follows: *p ≤ 0.05; **p ≤ 0.01; ***p ≤ 0.001; ****p < 0.0001.

## Acknowledgements

SNS was supported by a PhD scholarship at UCL funded by the Ministry of Education of Saudi Arabia and King Abdulaziz University. This work was also funded by the Medical Research Council (MRC) grant MR/W001241/1 to AF. NAB was supported by a PhD scholarship at UCL funded by the Government of Oman and AR was supported by a PhD scholarship at UCL funded by the Ministry of Education of Saudi Arabia and Prince Sattam Bin Abdulaziz University. We thank Sooyeon Chang, Adriana Albuquerque, Peter Steinberger, Ursula Demael, Hans Stauss for technical support and reagents, and Benedict Seddon and Clare Jolly for advice and helpful discussions. We are grateful to Petronela Ancuta for reading and commenting the manuscript. We thank the UCL Flow Cytometry Core Facility team Janani Sivakumaran Nguyen and Sam Blanchett for excellent support.

## Authors’ contributions

SNS, NAB designed and performed experiments, analysed the data and reviewed the manuscript, AR analysed the data and reviewed the manuscript, AF designed experiments, analysed the data and wrote the paper.

## Supporting information

**Fig. S1. Optimisation of LRAs to reactivate HIV-1 in latently infected Jurkat cells.** Latently infected Jurkat cells were activated for 24 hours (48 hours for the FOXO-1 inhibitor) with five different concentrations of (A) PMA, (B) PHA, (C) TLR7 agonist or TLR8 agonists, (D) Ionomycin, and (E) the FOXO-1 inhibitor. The cells were then analysed by flow cytometry to measure, from left to right, the percentage of GFP+ cells, GFP MFI, the percentage of CD69+ or CD71+ cells and cell viability using the same gating strategy shown in Fig 1 E. Bar graphs show the average values ± SD (n = 6 for GFP) and (n = 3 for viability, CD69, and CD71). Significance was calculated using a one-way ANOVA with Dunnett’s correction. *=p≤0.05; **=p≤0.01; ***=p≤0.001; ****=p<0.0001.

**Fig. S2. Optimisation of cytokines to reactivate HIV-1 in latently infected Jurkat cells.** (A) Latently infected Jurkat cells were activated for 24 hours with different concentrations of TNF-α and analysed by flow cytometry to measure, from left to right, the percentage of GFP+ cells, GFP MFI, the percentage of CD69+ cells and cell viability using the same gating strategy shown in Fig 1 E, n=6 for GFP and n = 3 for viability and activation markers. (B) latently infected cells were stimulated with different concentrations of IL-2, IL-6, IL-7+IL-15, IL-7 alone or IL-15 alone and analysed by flow cytometry to measure the percentage of GFP+ cells and (C) the GFP MFI (right panel). Bar graphs show the average values ± SD, n = 3. (D) Cell toxicity was analysed by flow cytometry using forward vs. side-scatter profiles. (E) Jurkat cells and primary CD4+ T cells were stained for CD127, the IL-7 and IL-15 receptor. A representative flow cytometry plot showing the percentage of CD127+ cells in CD4+ T cells.

**Fig. S3. AUY922 represses HIV-1 reactivation in latently infected Jurkat cells with no detectable cell toxicity.** Latently Jurkat cells were treated with a fixed concentration of LRAs alone or with different concentrations of AUY922 for 24 hours, except for FOXO-1, which was incubated for 48 hours. Cells were then stained with live or dead blue stain and analysed by flow cytometry to assess cell toxicity.

**Fig. S4. Gating strategy and FMO**. (A) Primary CD4+ T cells were analysed by spectral flow cytometry and positive gates were established by staining with the 18- antibody panel minus one (fluorescence minus one or FMO). (B) Representative flow cytometry plots and gating strategy of T cell subsets from one donor gated from live CD3+CD4+ cells.

**Fig. S5. The effect of AUY922 treatment on different CD4+ T cell populations**. CD4+ T cells were isolated from PBMCs and treated with IL-2 only (No stim) or anti- CD3/CD28 Abs + IL-2 for 72 hours, and AUY922 [25 nM] or DMSO added 48 hours post stimulation. Cells were analysed by flow cytometry 24 hours after the addition of AUY922. tSNE data of the T subsets, activation, and inhibitory markers were generated by FlowJo. A) tSNE plot for donor 2. B) tSNE plot for donor 3. C) tSNE plot for donor 4.

**Fig. S6. The effect of AUY922 treatment on different cytokine production.** Supernatant from latently infected CD4+ T cells, which were re-stimulated with anti- CD3/CD28 Abs in the presence or absence of DMSO (control), AUY922 (25 nM), or AUY922 (50 nM), was collected 48 hours after re-stimulation and used to measure different cytokines concentration. (A) IFN-γ, (B) IL-4, (C) IL-17A, (D) IL-10, and (E) TNF-α. Data are presented as mean ± standard error of the mean (SEM) n=5. Concentrations are shown in pg/ml. Statistical significance was determined using a Paired two-tailed Student’s t-test. *P < 0.05, **P < 0.01, ***P < 0.001. Pairwise comparisons were: No stim-IL-2 and anti-CD3/CD28; anti-CD3/CD28 and AUY922 25nM; anti-CD3/CD28 and AUY922 50 nM.

**S1 Table.** LRAs used to reactivate HIV-1 in latently infected Jurkat cells.

**S2 Table**. Concentrations of LRAs used to reactivate latent HIV-1 in Jurkat T cells.

**S3 Table**. T cell markers used for flow cytometry.

**S4 Table**. Markers used to identify T cell subsets, activation and inhibitory marker

## References

1. Simon V, Ho DD, Abdool Karim Q. HIV/AIDS epidemiology, pathogenesis, prevention, and treatment. Lancet. 2006;368(9534):489-504. doi: 10.1016/S0140-6736(06)69157-5. PubMed PMID: 16890836; PubMed Central PMCID: PMCPMC2913538.

2. Heaton RK, Clifford DB, Franklin DR, Jr., Woods SP, Ake C, Vaida F, et al. HIV-associated neurocognitive disorders persist in the era of potent antiretroviral therapy: CHARTER Study. Neurology. 2010;75(23):2087–96. doi: 10.1212/WNL.0b013e318200d727. PubMed PMID: 21135382; PubMed Central PMCID: PMCPMC2995535.

3. Trickey A, Sabin CA, Burkholder G, Crane H, d’Arminio Monforte A, Egger M, et al. Life expectancy after 2015 of adults with HIV on long-term antiretroviral therapy in Europe and North America: a collaborative analysis of cohort studies. Lancet HIV. 2023;10(5):e295-e307. Epub 20230320. doi: 10.1016/S2352-3018(23)00028-0. PubMed PMID: 36958365; PubMed Central PMCID: PMCPMC10288029.

4. Colby DJ, Trautmann L, Pinyakorn S, Leyre L, Pagliuzza A, Kroon E, et al. Rapid HIV RNA rebound after antiretroviral treatment interruption in persons durably suppressed in Fiebig I acute HIV infection. Nat Med. 2018;24(7):923–6. Epub 20180611. doi: 10.1038/s41591-018-0026-6. PubMed PMID: 29892063; PubMed Central PMCID: PMCPMC6092240.

5. Cohn LB, Chomont N, Deeks SG. The Biology of the HIV-1 Latent Reservoir and Implications for Cure Strategies. Cell Host Microbe. 2020;27(4):519–30. doi: 10.1016/j.chom.2020.03.014. PubMed PMID: 32272077; PubMed Central PMCID: PMCPMC7219958.

6. Veenhuis RT, Abreu CM, Shirk EN, Gama L, Clements JE. HIV replication and latency in monocytes and macrophages. Semin Immunol. 2021;51:101472. Epub 20210227. doi: 10.1016/j.smim.2021.101472. PubMed PMID: 33648815; PubMed Central PMCID: PMCPMC10171083.

7. Armani-Tourret M, Bone B, Tan TS, Sun W, Bellefroid M, Struyve T, et al. Immune targeting of HIV-1 reservoir cells: a path to elimination strategies and cure. Nat Rev Microbiol. 2024. Epub 20240209. doi: 10.1038/s41579-024-01010-8. PubMed PMID: 38337034.

8. Utay NS, Hunt PW. Role of immune activation in progression to AIDS. Curr Opin HIV AIDS. 2016;11(2):131–7. doi: 10.1097/COH.0000000000000242. PubMed PMID: 26731430; PubMed Central PMCID: PMCPMC4750472.

9. Schouten J, Wit FW, Stolte IG, Kootstra NA, van der Valk M, Geerlings SE, et al. Cross-sectional comparison of the prevalence of age-associated comorbidities and their risk factors between HIV-infected and uninfected individuals: the AGEhIV cohort study. Clin Infect Dis. 2014;59(12):1787–97. Epub 20140902. doi: 10.1093/cid/ciu701. PubMed PMID: 25182245.

10. Buck AM, LaFranchi BH, Henrich TJ. Gaining momentum: stem cell therapies for HIV cure. Curr Opin HIV AIDS. 2024. Epub 20240426. doi: 10.1097/COH.0000000000000859. PubMed PMID: 38686850.

11. Siliciano JD, Siliciano RF. HIV cure: The daunting scale of the problem. Science. 2024;383(6684):703-5. Epub 20240215. doi: 10.1126/science.adk1831. PubMed PMID: 38359111.

12. Pereira LA, Bentley K, Peeters A, Churchill MJ, Deacon NJ. A compilation of cellular transcription factor interactions with the HIV-1 LTR promoter. Nucleic Acids Res. 2000;28(3):663–8. doi: 10.1093/nar/28.3.663. PubMed PMID: 10637316; PubMed Central PMCID: PMCPMC102541.

13. Anderson I, Low JS, Weston S, Weinberger M, Zhyvoloup A, Labokha AA, et al. Heat shock protein 90 controls HIV-1 reactivation from latency. Proc Natl Acad Sci U S A. 2014;111(15):E1528-37. Epub 20140331. doi: 10.1073/pnas.1320178111. PubMed PMID: 24706778; PubMed Central PMCID: PMCPMC3992654.

14. Bell B, Sadowski I. Ras-responsiveness of the HIV-1 LTR requires RBF-1 and RBF-2 binding sites. Oncogene. 1996;13(12):2687–97. PubMed PMID: 9000143.

15. Brooks DG, Arlen PA, Gao L, Kitchen CM, Zack JA. Identification of T cell- signaling pathways that stimulate latent HIV in primary cells. Proc Natl Acad Sci U S A. 2003;100(22):12955–60. Epub 20031020. doi: 10.1073/pnas.2233345100. PubMed PMID: 14569007; PubMed Central PMCID: PMCPMC240726.

16. Chun TW, Engel D, Mizell SB, Ehler LA, Fauci AS. Induction of HIV-1 replication in latently infected CD4+ T cells using a combination of cytokines. J Exp Med. 1998;188(1):83–91. doi: 10.1084/jem.188.1.83. PubMed PMID: 9653086; PubMed Central PMCID: PMCPMC2525548.

17. Dobrowolski C, Valadkhan S, Graham AC, Shukla M, Ciuffi A, Telenti A, et al. Entry of Polarized Effector Cells into Quiescence Forces HIV Latency. mBio. 2019;10(2). Epub 20190326. doi: 10.1128/mBio.00337-19. PubMed PMID: 30914509; PubMed Central PMCID: PMCPMC6437053.

18. Van Lint C, Bouchat S, Marcello A. HIV-1 transcription and latency: an update. Retrovirology. 2013;10:67. Epub 20130626. doi: 10.1186/1742-4690-10-67. PubMed PMID: 23803414; PubMed Central PMCID: PMCPMC3699421.

19. Mbonye U, Karn J. The cell biology of HIV-1 latency and rebound. Retrovirology. 2024;21(1):6. Epub 20240405. doi: 10.1186/s12977-024-00639-w. PubMed PMID: 38580979; PubMed Central PMCID: PMCPMC10996279.

20. Mbonye U, Leskov K, Shukla M, Valadkhan S, Karn J. Biogenesis of P-TEFb in CD4+ T cells to reverse HIV latency is mediated by protein kinase C (PKC)- independent signaling pathways. PLoS Pathog. 2021;17(9):e1009581. Epub 20210916. doi: 10.1371/journal.ppat.1009581. PubMed PMID: 34529720; PubMed Central PMCID: PMCPMC8478230.

21. Chatterjee D, Zhang Y, Ngassaki-Yoka CD, Dutilleul A, Khalfi S, Hernalsteens O, et al. Identification of aryl hydrocarbon receptor as a barrier to HIV-1 infection and outgrowth in CD4(+) T cells. Cell Rep. 2023;42(6):112634. Epub 20230612. doi: 10.1016/j.celrep.2023.112634. PubMed PMID: 37310858; PubMed Central PMCID: PMCPMC10592455.

22. Wiche Salinas TR, Zhang Y, Sarnello D, Zhyvoloup A, Marchand LR, Fert A, et al. Th17 cell master transcription factor RORC2 regulates HIV-1 gene expression and viral outgrowth. Proc Natl Acad Sci U S A. 2021;118(48). doi: 10.1073/pnas.2105927118. PubMed PMID: 34819367; PubMed Central PMCID: PMCPMC8640723.

23. Li C, Mori L, Valente ST. The Block-and-Lock Strategy for Human Immunodeficiency Virus Cure: Lessons Learned from Didehydro-Cortistatin A. J Infect Dis. 2021;223(12 Suppl 2):46-53. doi: 10.1093/infdis/jiaa681. PubMed PMID: 33586776; PubMed Central PMCID: PMCPMC7883021.

24. Li C, Mousseau G, Valente ST. Tat inhibition by didehydro-Cortistatin A promotes heterochromatin formation at the HIV-1 long terminal repeat. Epigenetics Chromatin. 2019;12(1):23. Epub 20190416. doi: 10.1186/s13072-019-0267-8. PubMed PMID: 30992052; PubMed Central PMCID: PMCPMC6466689.

25. Mediouni S, Chinthalapudi K, Ekka MK, Usui I, Jablonski JA, Clementz MA, et al. Didehydro-Cortistatin A Inhibits HIV-1 by Specifically Binding to the Unstructured Basic Region of Tat. mBio. 2019;10(1). Epub 20190205. doi: 10.1128/mBio.02662-18. PubMed PMID: 30723126; PubMed Central PMCID: PMCPMC6368365.

26. Vozzolo L, Loh B, Gane PJ, Tribak M, Zhou L, Anderson I, et al. Gyrase B inhibitor impairs HIV-1 replication by targeting Hsp90 and the capsid protein. J Biol Chem. 2010;285(50):39314–28. Epub 20101011. doi: 10.1074/jbc.M110.155275. PubMed PMID: 20937817; PubMed Central PMCID: PMCPMC2998086.

27. Roesch F, Meziane O, Kula A, Nisole S, Porrot F, Anderson I, et al. Hyperthermia stimulates HIV-1 replication. PLoS Pathog. 2012;8(7):e1002792. Epub 20120712. doi: 10.1371/journal.ppat.1002792. PubMed PMID: 22807676; PubMed Central PMCID: PMCPMC3395604.

28. Joshi P, Maidji E, Stoddart CA. Inhibition of Heat Shock Protein 90 Prevents HIV Rebound. J Biol Chem. 2016;291(19):10332–46. Epub 20160308. doi: 10.1074/jbc.M116.717538. PubMed PMID: 26957545; PubMed Central PMCID: PMCPMC4858980.

29. Pan XY, Zhao W, Zeng XY, Lin J, Li MM, Shen XT, et al. Heat Shock Factor 1 Mediates Latent HIV Reactivation. Sci Rep. 2016;6:26294. Epub 20160518. doi: 10.1038/srep26294. PubMed PMID: 27189267; PubMed Central PMCID: PMCPMC4870680.

30. Low JS, Fassati A. Hsp90: a chaperone for HIV-1. Parasitology. 2014;141(9):1192–202. doi: 10.1017/S0031182014000298. PubMed PMID: 25004926.

31. Painter MM, Zaikos TD, Collins KL. Quiescence Promotes Latent HIV Infection and Resistance to Reactivation from Latency with Histone Deacetylase Inhibitors. J Virol. 2017;91(24). Epub 20171130. doi: 10.1128/JVI.01080-17. PubMed PMID: 29021396; PubMed Central PMCID: PMCPMC5709582.

32. Pan XY, Zhao W, Wang CY, Lin J, Zeng XY, Ren RX, et al. Heat Shock Protein 90 Facilitates Latent HIV Reactivation through Maintaining the Function of Positive Transcriptional Elongation Factor b (p-TEFb) under Proteasome Inhibition. J Biol Chem. 2016;291(50):26177–87. Epub 20161031. doi: 10.1074/jbc.M116.743906. PubMed PMID: 27799305; PubMed Central PMCID: PMCPMC5207085.

33. Peng W, Hong Z, Chen X, Gao H, Dai Z, Zhao J, et al. Thiostrepton Reactivates Latent HIV-1 through the p-TEFb and NF-kappaB Pathways Mediated by Heat Shock Response. Antimicrob Agents Chemother. 2020;64(5). Epub 20200421. doi: 10.1128/AAC.02328-19. PubMed PMID: 32094131; PubMed Central PMCID: PMCPMC7179580.

34. O’Keeffe B, Fong Y, Chen D, Zhou S, Zhou Q. Requirement for a kinase- specific chaperone pathway in the production of a Cdk9/cyclin T1 heterodimer responsible for P-TEFb-mediated tat stimulation of HIV-1 transcription. J Biol Chem. 2000;275(1):279–87. doi: 10.1074/jbc.275.1.279. PubMed PMID: 10617616.

35. Li ZN, Luo Y. HSP90 inhibitors and cancer: Prospects for use in targeted therapies (Review). Oncol Rep. 2023;49(1). Epub 20221111. doi: 10.3892/or.2022.8443. PubMed PMID: 36367182; PubMed Central PMCID: PMCPMC9685368.

36. Sessa C, Shapiro GI, Bhalla KN, Britten C, Jacks KS, Mita M, et al. First-in- human phase I dose-escalation study of the HSP90 inhibitor AUY922 in patients with advanced solid tumors. Clin Cancer Res. 2013;19(13):3671–80. Epub 20130611. doi: 10.1158/1078-0432.CCR-12-3404. PubMed PMID: 23757357.

37. Taipale M, Jarosz DF, Lindquist S. HSP90 at the hub of protein homeostasis: emerging mechanistic insights. Nat Rev Mol Cell Biol. 2010;11(7):515–28. Epub 20100609. doi: 10.1038/nrm2918. PubMed PMID: 20531426.

38. Boulon S, Pradet-Balade B, Verheggen C, Molle D, Boireau S, Georgieva M, et al. HSP90 and its R2TP/Prefoldin-like cochaperone are involved in the cytoplasmic assembly of RNA polymerase II. Mol Cell. 2010;39(6):912–24. doi: 10.1016/j.molcel.2010.08.023. PubMed PMID: 20864038; PubMed Central PMCID: PMCPMC4333224.

39. Jordan A, Bisgrove D, Verdin E. HIV reproducibly establishes a latent infection after acute infection of T cells in vitro. EMBO J. 2003;22(8):1868–77. doi: 10.1093/emboj/cdg188. PubMed PMID: 12682019; PubMed Central PMCID: PMCPMC154479.

40. Spina CA, Anderson J, Archin NM, Bosque A, Chan J, Famiglietti M, et al. An in-depth comparison of latent HIV-1 reactivation in multiple cell model systems and resting CD4+ T cells from aviremic patients. PLoS Pathog. 2013;9(12):e1003834. Epub 20131226. doi: 10.1371/journal.ppat.1003834. PubMed PMID: 24385908; PubMed Central PMCID: PMCPMC3873446.

41. Saleh S, Wightman F, Ramanayake S, Alexander M, Kumar N, Khoury G, et al. Expression and reactivation of HIV in a chemokine induced model of HIV latency in primary resting CD4+ T cells. Retrovirology. 2011;8:80. Epub 20111012. doi: 10.1186/1742-4690-8-80. PubMed PMID: 21992606; PubMed Central PMCID: PMCPMC3215964.

42. Covino DA, Desimio MG, Doria M. Impact of IL-15 and latency reversing agent combinations in the reactivation and NK cell-mediated suppression of the HIV reservoir. Sci Rep. 2022;12(1):18567. Epub 20221103. doi: 10.1038/s41598-022-23010-5. PubMed PMID: 36329160; PubMed Central PMCID: PMCPMC9633760.

43. Chen HC, Martinez JP, Zorita E, Meyerhans A, Filion GJ. Position effects influence HIV latency reversal. Nat Struct Mol Biol. 2017;24(1):47–54. Epub 20161121. doi: 10.1038/nsmb.3328. PubMed PMID: 27870832.

44. Vansant G, Chen HC, Zorita E, Trejbalova K, Miklik D, Filion G, et al. The chromatin landscape at the HIV-1 provirus integration site determines viral expression. Nucleic Acids Res. 2020;48(14):7801–17. doi: 10.1093/nar/gkaa536. PubMed PMID: 32597987; PubMed Central PMCID: PMCPMC7641320.

45. Zhyvoloup A, Melamed A, Anderson I, Planas D, Lee CH, Kriston-Vizi J, et al. Digoxin reveals a functional connection between HIV-1 integration preference and T- cell activation. PLoS Pathog. 2017;13(7):e1006460. Epub 20170720. doi: 10.1371/journal.ppat.1006460. PubMed PMID: 28727807; PubMed Central PMCID: PMCPMC5519191.

46. Yang HC, Xing S, Shan L, O’Connell K, Dinoso J, Shen A, et al. Small-molecule screening using a human primary cell model of HIV latency identifies compounds that reverse latency without cellular activation. J Clin Invest. 2009;119(11):3473–86. Epub 20091001. doi: 10.1172/JCI39199. PubMed PMID: 19805909; PubMed Central PMCID: PMCPMC2769176.

47. Bosque A, Planelles V. Induction of HIV-1 latency and reactivation in primary memory CD4+ T cells. Blood. 2009;113(1):58–65. Epub 20081010. doi: 10.1182/blood-2008-07-168393. PubMed PMID: 18849485; PubMed Central PMCID: PMCPMC2614643.

48. Meas HZ, Haug M, Beckwith MS, Louet C, Ryan L, Hu Z, et al. Sensing of HIV-1 by TLR8 activates human T cells and reverses latency. Nat Commun. 2020;11(1):147. Epub 20200109. doi: 10.1038/s41467-019-13837-4. PubMed PMID: 31919342; PubMed Central PMCID: PMCPMC6952430.

49. Simms PE, Ellis TM. Utility of flow cytometric detection of CD69 expression as a rapid method for determining poly- and oligoclonal lymphocyte activation. Clin Diagn Lab Immunol. 1996;3(3):301–4. doi: 10.1128/cdli.3.3.301-304.1996. PubMed PMID: 8705673; PubMed Central PMCID: PMCPMC170336.

50. Tsai A, Irrinki A, Kaur J, Cihlar T, Kukolj G, Sloan DD, et al. Toll-Like Receptor 7 Agonist GS-9620 Induces HIV Expression and HIV-Specific Immunity in Cells from HIV-Infected Individuals on Suppressive Antiretroviral Therapy. J Virol. 2017;91(8). Epub 20170329. doi: 10.1128/JVI.02166-16. PubMed PMID: 28179531; PubMed Central PMCID: PMCPMC5375698.

51. Vallejo-Gracia A, Chen IP, Perrone R, Besnard E, Boehm D, Battivelli E, et al. FOXO1 promotes HIV latency by suppressing ER stress in T cells. Nat Microbiol. 2020;5(9):1144–57. Epub 20200615. doi: 10.1038/s41564-020-0742-9. PubMed PMID: 32541947; PubMed Central PMCID: PMCPMC7483895.

52. Roux A, Leroy H, De Muylder B, Bracq L, Oussous S, Dusanter-Fourt I, et al. FOXO1 transcription factor plays a key role in T cell-HIV-1 interaction. PLoS Pathog. 2019;15(5):e1007669. Epub 20190501. doi: 10.1371/journal.ppat.1007669. PubMed PMID: 31042779; PubMed Central PMCID: PMCPMC6513100.

53. Eccles SA, Massey A, Raynaud FI, Sharp SY, Box G, Valenti M, et al. NVP- AUY922: a novel heat shock protein 90 inhibitor active against xenograft tumor growth, angiogenesis, and metastasis. Cancer Res. 2008;68(8):2850–60. doi: 10.1158/0008-5472.CAN-07-5256. PubMed PMID: 18413753.

54. Neckers L, Workman P. Hsp90 molecular chaperone inhibitors: are we there yet? Clin Cancer Res. 2012;18(1):64–76. doi: 10.1158/1078-0432.CCR-11-1000. PubMed PMID: 22215907; PubMed Central PMCID: PMCPMC3252205.

55. Battivelli E, Dahabieh MS, Abdel-Mohsen M, Svensson JP, Tojal Da Silva I, Cohn LB, et al. Distinct chromatin functional states correlate with HIV latency reactivation in infected primary CD4(+) T cells. Elife. 2018;7. Epub 20180501. doi: 10.7554/eLife.34655. PubMed PMID: 29714165; PubMed Central PMCID: PMCPMC5973828.

56. Dar RD, Hosmane NN, Arkin MR, Siliciano RF, Weinberger LS. Screening for noise in gene expression identifies drug synergies. Science. 2014;344(6190):1392-6. Epub 20140605. doi: 10.1126/science.1250220. PubMed PMID: 24903562; PubMed Central PMCID: PMCPMC4122234.

57. Schulte TW, Neckers LM. The benzoquinone ansamycin 17-allylamino-17- demethoxygeldanamycin binds to HSP90 and shares important biologic activities with geldanamycin. Cancer Chemother Pharmacol. 1998;42(4):273–9. doi: 10.1007/s002800050817. PubMed PMID: 9744771.

58. Lee J, Mira-Arbibe L, Ulevitch RJ. TAK1 regulates multiple protein kinase cascades activated by bacterial lipopolysaccharide. Journal of leukocyte biology. 2000;68(6):909–15.

59. Ninomiya-Tsuji J, Kishimoto K, Hiyama A, Inoue J-i, Cao Z, Matsumoto K. The kinase TAK1 can activate the NIK-IκB as well as the MAP kinase cascade in the IL-1 signalling pathway. Nature. 1999;398(6724):252-6.

60. Shim J-H, Xiao C, Paschal AE, Bailey ST, Rao P, Hayden MS, et al. TAK1, but not TAB1 or TAB2, plays an essential role in multiple signaling pathways in vivo. Genes & development. 2005;19(22):2668–81.

61. Wang XD, Zhao CS, Wang QL, Zeng Q, Feng XZ, Li L, et al. The p38-interacting protein p38IP suppresses TCR and LPS signaling by targeting TAK1. EMBO reports. 2020;21(7):e48035.

62. Sun L, Deng L, Ea CK, Xia ZP, Chen ZJ. The TRAF6 ubiquitin ligase and TAK1 kinase mediate IKK activation by BCL10 and MALT1 in T lymphocytes. Mol Cell. 2004;14(3):289–301. doi: 10.1016/s1097-2765(04)00236-9. PubMed PMID: 15125833.

63. Hwang JR, Byeon Y, Kim D, Park SG. Recent insights of T cell receptor- mediated signaling pathways for T cell activation and development. Exp Mol Med. 2020;52(5):750–61. Epub 20200521. doi: 10.1038/s12276-020-0435-8. PubMed PMID: 32439954; PubMed Central PMCID: PMCPMC7272404.

64. Zhu L, Lama S, Tu L, Dusting GJ, Wang JH, Liu GS. TAK1 signaling is a potential therapeutic target for pathological angiogenesis. Angiogenesis. 2021;24(3):453–70. Epub 20210510. doi: 10.1007/s10456-021-09787-5. PubMed PMID: 33973075.

65. Shi L, Zhang Z, Fang S, Xu J, Liu J, Shen J, et al. Heat shock protein 90 (Hsp90) regulates the stability of transforming growth factor β-activated kinase 1 (TAK1) in interleukin-1β-induced cell signaling. Molecular immunology. 2009;46(4):541–50.

66. Liu XY, Seh CC, Cheung PC. HSP90 is required for TAK1 stability but not for its activation in the pro-inflammatory signaling pathway. FEBS letters. 2008;582(29):4023–31.

67. Liu R, Lin Y, Jia R, Geng Y, Liang C, Tan J, et al. HIV-1 Vpr stimulates NF- kappaB and AP-1 signaling by activating TAK1. Retrovirology. 2014;11:45. Epub 20140609. doi: 10.1186/1742-4690-11-45. PubMed PMID: 24912525; PubMed Central PMCID: PMCPMC4057933.

68. Hwang J-R, Byeon Y, Kim D, Park S-G. Recent insights of T cell receptor- mediated signaling pathways for T cell activation and development. Experimental & molecular medicine. 2020;52(5):750–61.

69. Shah K, Al-Haidari A, Sun J, Kazi JU. T cell receptor (TCR) signaling in health and disease. Signal transduction and targeted therapy. 2021;6(1):412.

70. Wu J, Powell F, Larsen NA, Lai Z, Byth KF, Read J, et al. Mechanism and in vitro pharmacology of TAK1 inhibition by (5Z)-7-Oxozeaenol. ACS Chem Biol. 2013;8(3):643–50. Epub 20130107. doi: 10.1021/cb3005897. PubMed PMID: 23272696.

71. Jutz S, Leitner J, Schmetterer K, Doel-Perez I, Majdic O, Grabmeier- Pfistershammer K, et al. Assessment of costimulation and coinhibition in a triple parameter T cell reporter line: Simultaneous measurement of NF-kappaB, NFAT and AP-1. J Immunol Methods. 2016;430:10-20. Epub 20160115. doi: 10.1016/j.jim.2016.01.007. PubMed PMID: 26780292.

72. Mousset CM, Hobo W, Woestenenk R, Preijers F, Dolstra H, van der Waart AB. Comprehensive Phenotyping of T Cells Using Flow Cytometry. Cytometry A. 2019;95(6):647–54. Epub 20190204. doi: 10.1002/cyto.a.23724. PubMed PMID: 30714682.

73. van den Broek T, Borghans JAM, van Wijk F. The full spectrum of human naive T cells. Nat Rev Immunol. 2018;18(6):363–73. doi: 10.1038/s41577-018-0001-y. PubMed PMID: 29520044.

74. Kumar BV, Connors TJ, Farber DL. Human T Cell Development, Localization, and Function throughout Life. Immunity. 2018;48(2):202-13. doi: 10.1016/j.immuni.2018.01.007. PubMed PMID: 29466753; PubMed Central PMCID: PMCPMC5826622.

75. Jameson SC, Masopust D. Understanding Subset Diversity in T Cell Memory. Immunity. 2018;48(2):214–26. doi: 10.1016/j.immuni.2018.02.010. PubMed PMID: 29466754; PubMed Central PMCID: PMCPMC5863745.

76. Golubovskaya V, Wu L. Different Subsets of T Cells, Memory, Effector Functions, and CAR-T Immunotherapy. Cancers (Basel). 2016;8(3). Epub 20160315. doi: 10.3390/cancers8030036. PubMed PMID: 26999211; PubMed Central PMCID: PMCPMC4810120.

77. Raphael I, Nalawade S, Eagar TN, Forsthuber TG. T cell subsets and their signature cytokines in autoimmune and inflammatory diseases. Cytokine. 2015;74(1):5–17. Epub 20141030. doi: 10.1016/j.cyto.2014.09.011. PubMed PMID: 25458968; PubMed Central PMCID: PMCPMC4416069.

78. Maecker HT, McCoy JP, Nussenblatt R. Standardizing immunophenotyping for the Human Immunology Project. Nat Rev Immunol. 2012;12(3):191–200. Epub 20120217. doi: 10.1038/nri3158. PubMed PMID: 22343568; PubMed Central PMCID: PMCPMC3409649.

79. Aspalter RM, Eibl MM, Wolf HM. Regulation of TCR-mediated T cell activation by TNF-RII. Journal of Leucocyte Biology. 2003;74(4):572–82.

80. Fenwick C, Joo V, Jacquier P, Noto A, Banga R, Perreau M, et al. T-cell exhaustion in HIV infection. Immunological reviews. 2019;292(1):149–63.

81. Pardons M, Baxter AE, Massanella M, Pagliuzza A, Fromentin R, Dufour C, et al. Single-cell characterization and quantification of translation-competent viral reservoirs in treated and untreated HIV infection. PLoS Pathog. 2019;15(2):e1007619. Epub 20190227. doi: 10.1371/journal.ppat.1007619. PubMed PMID: 30811499; PubMed Central PMCID: PMCPMC6411230.

82. Xie G, Luo X, Ma T, Frouard J, Neidleman J, Hoh R, et al. Characterization of HIV-induced remodeling reveals differences in infection susceptibility of memory CD4(+) T cell subsets in vivo. Cell Rep. 2021;35(4):109038. doi: 10.1016/j.celrep.2021.109038. PubMed PMID: 33910003; PubMed Central PMCID: PMCPMC8202093.

83. Van der Maaten L, Hinton G. Visualizing data using t-SNE. Journal of machine learning research. 2008;9(11).

84. Zhou L, Sokolskaja E, Jolly C, James W, Cowley SA, Fassati A. Transportin 3 promotes a nuclear maturation step required for efficient HIV-1 integration. PLoS Pathog. 2011;7(8):e1002194. Epub 20110825. doi: 10.1371/journal.ppat.1002194. PubMed PMID: 21901095; PubMed Central PMCID: PMCPMC3161976.

85. Geginat J, Lanzavecchia A, Sallusto F. Proliferation and differentiation potential of human CD8+ memory T-cell subsets in response to antigen or homeostatic cytokines. Blood. 2003;101(11):4260–6. Epub 20030206. doi: 10.1182/blood-2002-11-3577. PubMed PMID: 12576317.

86. Shipkova M, Wieland E. Surface markers of lymphocyte activation and markers of cell proliferation. Clinica chimica acta. 2012;413(17-18):1338–49.

87. Farber DL, Yudanin NA, Restifo NP. Human memory T cells: generation, compartmentalization and homeostasis. Nat Rev Immunol. 2014;14(1):24–35. Epub 20131213. doi: 10.1038/nri3567. PubMed PMID: 24336101; PubMed Central PMCID: PMCPMC4032067.

88. Kim YS, Alarcon SV, Lee S, Lee MJ, Giaccone G, Neckers L, et al. Update on Hsp90 inhibitors in clinical trial. Curr Top Med Chem. 2009;9(15):1479–92. doi: 10.2174/156802609789895728. PubMed PMID: 19860730; PubMed Central PMCID: PMCPMC7241864.

89. Schnaider T, Somogyi J, Csermely P, Szamel M. The Hsp90-specific inhibitor geldanamycin selectively disrupts kinase-mediated signaling events of T-lymphocyte activation. Cell stress & chaperones. 2000;5(1):52.

90. Hayashi K, Kamikawa Y. HSP90 is crucial for regulation of LAT expression in activated T cells. Molecular immunology. 2011;48(6-7):941–6.

91. Smith-Garvin JE, Koretzky GA, Jordan MS. T cell activation. Annu Rev Immunol. 2009;27:591-619. doi: 10.1146/annurev.immunol.021908.132706. PubMed PMID: 19132916; PubMed Central PMCID: PMCPMC2740335.

92. Bennett LD, Fox JM, Signoret N. Mechanisms regulating chemokine receptor activity. Immunology. 2011;134(3):246–56.

93. Sallusto F, Kremmer E, Palermo B, Hoy A, Ponath P, Qin S, et al. Switch in chemokine receptor expression upon TCR stimulation reveals novel homing potential for recently activated T cells. Eur J Immunol. 1999;29(6):2037–45. doi: 10.1002/(SICI)1521-4141(199906)29:06<2037::AID-IMMU2037>3.0.CO;2-V. PubMed PMID: 10382767.

94. Chiosis G, Neckers L. Tumor selectivity of Hsp90 inhibitors: the explanation remains elusive. ACS chemical biology. 2006;1(5):279–84.

95. Kamal A, Thao L, Sensintaffar J, Zhang L, Boehm MF, Fritz LC, et al. A high- affinity conformation of Hsp90 confers tumour selectivity on Hsp90 inhibitors. Nature. 2003;425(6956):407-10.

96. Unutmaz D, KewalRamani VN, Marmon S, Littman DR. Cytokine signals are sufficient for HIV-1 infection of resting human T lymphocytes. J Exp Med. 1999;189(11):1735–46. doi: 10.1084/jem.189.11.1735. PubMed PMID: 10359577; PubMed Central PMCID: PMCPMC2193071.

97. Ducrey-Rundquist O, Guyader M, Trono D. Modalities of interleukin-7-induced human immunodeficiency virus permissiveness in quiescent T lymphocytes. J Virol. 2002;76(18):9103–11. doi: 10.1128/jvi.76.18.9103-9111.2002. PubMed PMID:

98. 12186894; PubMed Central PMCID: PMCPMC136444.

98. Reuschl AK, Mesner D, Shivkumar M, Whelan MVX, Pallett LJ, Guerra- Assuncao JA, et al. HIV-1 Vpr drives a tissue residency-like phenotype during selective infection of resting memory T cells. Cell Rep. 2022;39(2):110650. doi: 10.1016/j.celrep.2022.110650.<otherinfo> PubMed PMID:</otherinfo> 35417711; PubMed Central PMCID: PMCPMC9350556.

99. Scripture-Adams DD, Brooks DG, Korin YD, Zack JA. Interleukin-7 induces expression of latent human immunodeficiency virus type 1 with minimal effects on T- cell phenotype. J Virol. 2002;76(24):13077–82. doi: 10.1128/jvi.76.24.13077-13082.2002. PubMed PMID: 12438635; PubMed Central PMCID: PMCPMC136703.

100. Wang FX, Xu Y, Sullivan J, Souder E, Argyris EG, Acheampong EA, et al. IL-7 is a potent and proviral strain-specific inducer of latent HIV-1 cellular reservoirs of infected individuals on virally suppressive HAART. J Clin Invest. 2005;115(1):128–37. doi: 10.1172/JCI22574. PubMed PMID: 15630452; PubMed Central PMCID: PMCPMC539197.

101. Mohammadi P, Di Iulio J, Munoz M, Martinez R, Bartha I, Cavassini M, et al. Dynamics of HIV latency and reactivation in a primary CD4+ T cell model. PLoS pathogens. 2014;10(5):e1004156.

102. Bosque A, Famiglietti M, Weyrich AS, Goulston C, Planelles V. Homeostatic proliferation fails to efficiently reactivate HIV-1 latently infected central memory CD4+ T cells. PLoS pathogens. 2011;7(10):e1002288.

103. Gattinoni L, Speiser DE, Lichterfeld M, Bonini C. T memory stem cells in health and disease. Nat Med. 2017;23(1):18–27. doi: 10.1038/nm.4241. PubMed PMID: 28060797; PubMed Central PMCID: PMCPMC6354775.

104. Bosque A, Planelles V. Induction of HIV-1 latency and reactivation in primary memory CD4+ T cells. Blood, The Journal of the American Society of Hematology. 2009;113(1):58–65.

105. Fromentin R, Chomont N, editors. HIV persistence in subsets of CD4+ T cells: 50 shades of reservoirs. Seminars in immunology; 2021: Elsevier.

106. Kulpa DA, Talla A, Brehm JH, Ribeiro SP, Yuan S, Bebin-Blackwell A-G, et al. Differentiation into an effector memory phenotype potentiates HIV-1 latency reversal in CD4+ T cells. Journal of virology. 2019;93(24):10.1128/jvi. 00969-19.

107. Grau-Exposito J, Luque-Ballesteros L, Navarro J, Curran A, Burgos J, Ribera E, et al. Latency reversal agents affect differently the latent reservoir present in distinct CD4+ T subpopulations. PLoS Pathog. 2019;15(8):e1007991. Epub 20190819. doi: 10.1371/journal.ppat.1007991. PubMed PMID: 31425551; PubMed Central PMCID: PMCPMC6715238.

108. Baxter AE, Niessl J, Fromentin R, Richard J, Porichis F, Charlebois R, et al. Single-Cell Characterization of Viral Translation-Competent Reservoirs in HIV- Infected Individuals. Cell Host Microbe. 2016;20(3):368–80. Epub 20160818. doi: 10.1016/j.chom.2016.07.015. PubMed PMID: 27545045; PubMed Central PMCID: PMCPMC5025389.

109. Magyar CTJ, Vashist YK, Stroka D, Kim-Fuchs C, Berger MD, Banz VM. Heat shock protein 90 (HSP90) inhibitors in gastrointestinal cancer: where do we currently stand?-A systematic review. J Cancer Res Clin Oncol. 2023;149(10):8039–50. Epub 20230326. doi: 10.1007/s00432-023-04689-z. PubMed PMID: 36966394; PubMed Central PMCID: PMCPMC10374781.

110. Lang JE, Forero-Torres A, Yee D, Yau C, Wolf D, Park J, et al. Safety and efficacy of HSP90 inhibitor ganetespib for neoadjuvant treatment of stage II/III breast cancer. NPJ Breast Cancer. 2022;8(1):128. Epub 20221201. doi: 10.1038/s41523-022-00493-z. PubMed PMID: 36456573; PubMed Central PMCID: PMCPMC9715670.

111. Geller R, Taguwa S, Frydman J. Broad action of Hsp90 as a host chaperone required for viral replication. Biochimica et Biophysica Acta (BBA)-Molecular Cell Research. 2012;1823(3):698–706.

112. Basha W, Kitagawa R, Uhara M, Imazu H, Uechi K, Tanaka J. Geldanamycin, a potent and specific inhibitor of Hsp90, inhibits gene expression and replication of human cytomegalovirus. Antiviral Chemistry and Chemotherapy. 2005;16(2):135–46.

113. Acchioni C, Sandini S, Acchioni M, Sgarbanti M. Co-Infections and Superinfections between HIV-1 and Other Human Viruses at the Cellular Level. Pathogens. 2024;13(5). Epub 20240424. doi: 10.3390/pathogens13050349. PubMed PMID: 38787201; PubMed Central PMCID: PMCPMC11124504.

114. Sauter D, Hotter D, Van Driessche B, Stürzel CM, Kluge SF, Wildum S, et al. Differential regulation of NF-κB-mediated proviral and antiviral host gene expression by primate lentiviral Nef and Vpu proteins. Cell reports. 2015;10(4):586–99.

115. Gummuluru S, Emerman M. Cell cycle- and Vpr-mediated regulation of human immunodeficiency virus type 1 expression in primary and transformed T-cell lines. J Virol. 1999;73(7):5422–30. doi: 10.1128/JVI.73.7.5422-5430.1999. PubMed PMID: 10364289; PubMed Central PMCID: PMCPMC112598.

116. Fassati A, Goff SP. Characterization of intracellular reverse transcription complexes of human immunodeficiency virus type 1. J Virol. 2001;75(8):3626–35. doi: 10.1128/JVI.75.8.3626-3635.2001. PubMed PMID: 11264352; PubMed Central PMCID: PMCPMC114854.

